# Ancestral reconstruction of sunflower karyotypes reveals non-random chromosomal evolution

**DOI:** 10.1101/737155

**Authors:** Kate L. Ostevik, Kieran Samuk, Loren H. Rieseberg

**Author notes:** Corresponding author: Kate Ostevik, Box 90338, 137 Biological Sciences, 130 Science Drive, Durham, NC, 27708.

## Abstract

Mapping the chromosomal rearrangements between species can inform our understanding of genome evolution, reproductive isolation, and speciation. Here we present a novel algorithm for identifying regions of synteny in pairs of genetic maps, which is implemented in the accompanying R package, syntR. The syntR algorithm performs as well as previous ad-hoc methods while being systematic, repeatable, and is applicable to mapping chromosomal rearrangements in any group of species. In addition, we present a systematic survey of chromosomal rearrangements in the annual sunflowers, which is a group known for extreme karyotypic diversity. We build high-density genetic maps for two subspecies of the prairie sunflower*, Helianthus petiolaris* ssp. *petiolaris* and *H. petiolaris* ssp. *fallax*.

Using syntR, and we identify blocks of synteny between these two subspecies and previously published high-density genetic maps. We reconstruct ancestral karyotypes for annual sunflowers using those synteny blocks and conservatively estimate that there have been 7.9 chromosomal rearrangements per million years – a high rate of chromosomal evolution. Although the rate of inversion is even higher than the rate of translocation in this group, we further find that every extant karyotype is distinguished by between 1 and 3 translocations involving only 8 of the 17 chromosomes. This non-random exchange suggests that specific chromosomes are prone to translocation and may thus contribute disproportionately to widespread hybrid sterility in sunflowers. These data deepen our understanding of chromosome evolution and confirm that *Helianthus* has an exceptional rate of chromosomal rearrangement that may facilitate similarly rapid diversification.

## Introduction

Organisms vary widely in the number and arrangement of their chromosomes – i.e., their karyotype. Interestingly, karyotypic differences are often associated with species boundaries and, therefore, suggest a link between chromosomal evolution and speciation (White 1978, King 1993). Indeed, it is well established that chromosomal rearrangements can contribute to reproductive isolation. Individuals heterozygous for divergent karyotypes are often sterile or inviable (King 1987, Lai et al. 2005, Stathos and Fishman 2014). Apart from directly causing hybrid sterility and inviability, chromosomal rearrangements can also facilitate the evolution of other reproductive barriers by extending genomic regions that are protected from introgression (Noor et al. 2001, Rieseberg 2001), accumulating genetic incompatibilities (Navarro and Barton 2003), and simplifying reinforcement (Trickett and Butlin 1994). Despite its prevalence and potentially important role in speciation, the general patterns of karyotypic divergence are still not well understood. Mapping and characterizing chromosomal rearrangements in many taxa is a critical step towards understanding their evolutionary dynamics.

The genus *Helianthus* (sunflowers) is well known to have particularly labile genome structure and is thus a viable system in which to map and characterize a variety of rearrangements. These sunflowers have several paleopolyploidy events in their evolutionary history (Barker et al. 2008, Barker et al. 2016, Badouin et al. 2017), have given rise to three homoploid hybrid species (Rieseberg 1991), and are prone to transposable element activity (Kawakami et al. 2011, Staton et al. 2012). Evidence in the form of hybrid pollen inviability, abnormal chromosome pairings during meiosis, and genetic map comparisons suggests that *Helianthus* karyotypes are unusually diverse (Heiser 1947, Heiser 1951, Heiser 1961, Whelan 1979, Chandler 1986, Rieseberg et al. 1995, Quillet et al. 1995, Burke et al. 2004, Heesacker et al. 2009, Barb et al. 2014). In fact, annual sunflowers have one of the highest described rates of chromosomal evolution across all plants and animals (Burke et al. 2004).

Studying chromosomal evolution within any group requires high-density genetic maps. Recently, Barb et al. (2014) built high-density genetic maps for the sunflower species *H. niveus* ssp. *tephrodes* and *H. argophyllus* and compared them to *H. annuus*. This analysis precisely mapped previously inferred karyotypes (Heiser 1951, Chandler 1986, Quillet et al. 1995), but only captured a small amount of the chromosomal variation in the annual sunflowers. For example, comparisons of genetic maps with limited marker density suggest that several chromosomal rearrangements differentiate *H. petiolaris* from *H. annuus* and (Rieseberg et al. 1995, Burke et al. 2004) and evidence from cytological surveys suggests that subspecies within *H. petiolaris* subspecies carry divergent karyotypes (Heiser 1961).

Adding high-density genetic maps of *H. petiolaris* subspecies to the Barb et al. (2014) analysis will allow us to: (1) precisely track additional rearrangements, (2) reconstruct ancestral karyotypes for the group, and (3) untangle overlapping rearrangements that can be obscured by directly comparing present-day karyotypes.

Another critical part of a multi-species comparative study of chromosome evolution using genetic map data is a systematic and repeatable method for identifying syntenic chromosomal regions (*sensu* Pevzner and Tesler 2003). These methods are especially important for cases with high marker density because breakpoints between synteny blocks can be blurred by mapping error, micro-rearrangements, and paralogy (Hackett and Broadfoot 2003, Choi et al. 2007, Barb et al. 2014, Bilton et al. 2018). In previous studies, synteny blocks have been found by a variety of ad-hoc methods, including counting all differences in marker order (Wu and Tanksley 2010), by visual inspection (Burke et al. 2004, Marone et al. 2012, Latta et al. 2019), or by manually applying simple rules like size thresholds (Heesacker et al. 2009, Barb et al. 2014, Rueppell et al. 2016) and Spearman’s rank comparisons (Berdan et al. 2014, Schlautman et al. 2017). However, these methods become intractable and prone to error when applied to very dense genetic maps. Furthermore, to our knowledge, there is no software available that identifies synteny blocks based on relative marker positions alone (i.e., without requiring reference genomes, sequence data, or markers with known orientations).

Here, with the goal of understanding chromosome evolution in *Helianthus* and more generally, we aimed to: (1) build high-density genetic maps for two subspecies of *Helianthus petiolaris*, (2) develop a method and software to systematically and repeatably identify synteny blocks from any number of paired genetic map positions, (3) reconstruct ancestral karyotypes for a subsection of annual sunflowers, and (4) detect general patterns of chromosomal rearrangement in *Helianthus*.

## Methods

### Study system

We focused on five closely related diploid (2*n* = 34) taxa from the annual clade of the genus *Helianthus* (Fig 1). These sunflowers are native to North America (Fig S1, Rogers et al. 1982) and are naturally self-incompatible (domesticated lineages of *H. annuus* are self-compatible). *Helianthus annuus* occurs throughout much of the central United States, often in somewhat heavy soils and along roadsides (Heiser 1947). *Helianthus petiolaris* occurs in sandier soils and is made up of two subspecies: *H. petiolaris* ssp. *petiolaris*, which is commonly found in the southern Great Plains, and *H. petiolaris* ssp. *fallax*, which is limited to more arid regions in Colorado, Utah, New Mexico, and Arizona (Heiser 1961). Where *H. petiolaris* and *H. annuus* are sympatric, gene flow occurs between the species (Strasburg and Rieseberg 2008). *Helianthus argophyllus* is primarily found along the east coast of Texas where it also overlaps and hybridizes with *H. annuus* (Baute et al. 2016). Finally, *H. niveus* ssp. *tephrodes* is a facultative perennial that grows in dunes from the southwestern US into Mexico.

**Figure 1.**
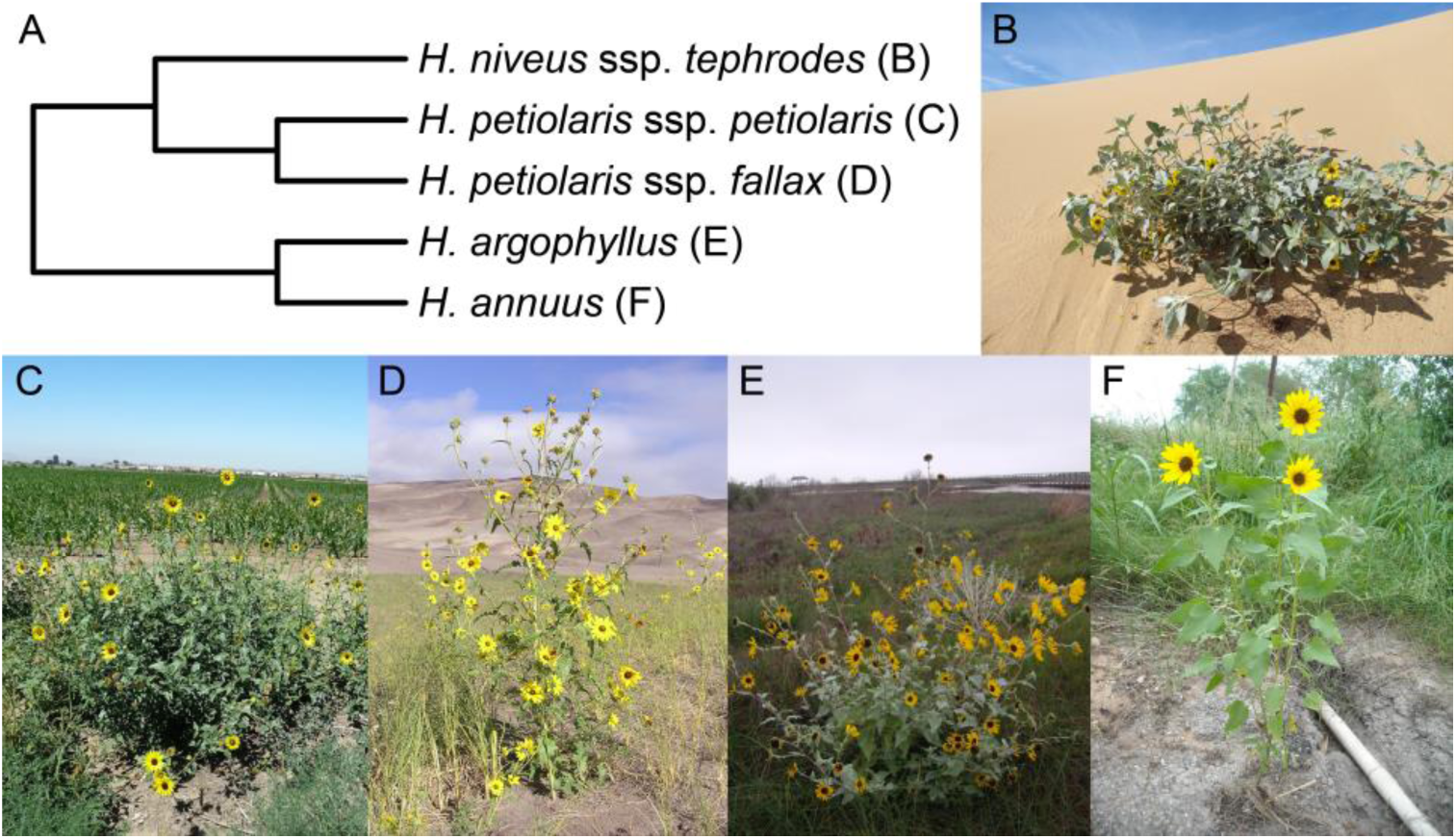
The sunflower taxa used in this study. A) Phylogenetic relationships based on Stephens et al. (2015) and Baute et al. (2016). B) *H. niveus* ssp. *tephrodes.* C) *H. petiolaris* ssp. *petiolaris.* D) *H. petiolaris* ssp. *fallax.* E) *argophyllus.* F) *H. annuus.* Photo credits: Brook Moyers (B, C, E & F) and Rose Andrew (D).

### Controlled crosses

To make genetic maps, we crossed an outbred individual with presumably high heterozygosity from each *H. petiolaris* subspecies to a homozygous inbred line of domesticated sunflower and genotyped the resulting F1 offspring. This test-cross design allows us to infer where recombination occurred in the heterozygous parents because we can reliably track the segregation of those parents’ alleles against a predictable background (Fig 2).

**Figure 2.**
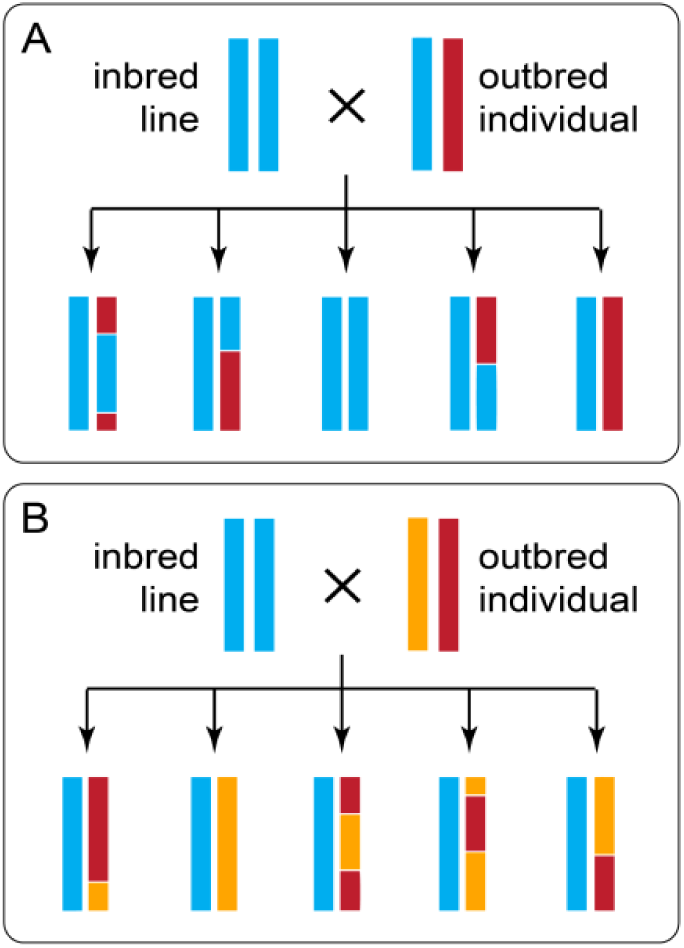
Diagram showing how a test-cross can be used to map the recombination events in an outbred individual that may (A) or may not (B) share alleles with the inbred line. Each line represents a chromosome, and the colors represent ancestry.

Specifically, we used pollen from a single *H. petiolaris ssp. petiolaris* plant (PI435836) and a single *H. petiolaris ssp. fallax* plant (PI435768) to fertilize individuals of a highly inbred and male sterile line of *H. annuus* (HA89cms). The self-incompatible *H. petiolaris* accessions were collected in central Colorado (PI435836, 39.741**°**, -105.342°, Boulder County) and the southeast corner of New Mexico (PI435768, 32.3**°**, -104.0°, Eddy County, Fig S1) and were maintained at large population sizes by the United States Department of Agriculture. When it was originally collected, accession PI435768 was classified *H. neglectus*. However, based on the location of the collection (Heiser 1961) and a more recent genetic analysis of the scale of differences between *H. petiolaris* ssp. *fallax* and *H. neglectus* (Raduski et al. 2010), we believe that this accession should be classified *H. petiolaris ssp. fallax*.

### Genotyping

We collected leaf tissue from 116 *H. annuus* x *H. petiolaris* ssp. *petiolaris* F1 seedlings and 132 *H. annuus* x *H. petiolaris* ssp. *fallax* F1 seedlings. We extracted DNA using a modified CTAB protocol (Doyle and Doyle 1987) and prepared individually barcoded genotyping-by-sequencing (GBS) libraries using a version the Poland et al. (2012) protocol. Our modified protocol includes steps to reduce the frequency of high-copy fragments (e.g., chloroplast and repetitive sequence) based on Shagina et al. (2010) and Matvienko et al. (2013) and steps to select specific fragment sizes for sequencing (see Ostevik 2016 appendix B for the full protocol).

Briefly, we digested 100ng of DNA from each individual with restriction enzymes (either *Pst*I-HF or *Pst*I-HF and *Msp*I) and ligated individual barcodes and common adapters to the digested DNA. We pooled barcoded fragments from up to 192 individuals, cleaned and concentrated the libraries using SeraMag Speed Beads made in-house (Rohland and Reich 2012), and amplified fragments using 12 cycles of PCR. We depleted high-copy fragments based on Todesco et al. (2019) using the following steps: (1) denature the libraries using high temperatures, (2) allow the fragments to re-hybridize, (3) digest the double-stranded fragments with duplex specific nuclease (Zhulidov et al. 2004), and (4) amplify the undigested fragments using another 12 cycles of PCR. We ran the libraries out on a 1.5% agarose gel and extracted 300-800 bp fragments using a Zymoclean Gel DNA Recovery kit (Zymo Research, Irvine, USA). Then, following additional library cleanup and quality assessment, we sequenced paired-ends of our libraries on an Illumina HiSeq 2000 (Illumina Inc., San Diego, CA, USA).

To call variants, we used a pipeline that combines the Burrows-Wheeler Aligner version 0.7.15 (BWA, Li & Durbin 2010) and the Genome Analysis Toolkit version 3.7 (GATK, McKenna et al. 2010). First, we demultiplexed the data using sabre (https://github.com/najoshi/sabre, Accessed 27 Jan 2017). Next, we aligned reads to the *H. annuus* reference (HanXRQr1.0-20151230, Badouin et al. 2017) with ‘bwa-mem’ (Li 2013), called variants with GATK ‘HaplotypeCaller’, and jointly genotyped all samples within a cross type with GATK ‘GentypeGVCFs’. We split variants into SNPs and indels and filtered each marker type using hard-filtration criteria suggested in the GATK best practices (DePristo et al. 2011, Van der Auwera et al. 2013). Specifically, we removed SNPs that had quality by depth scores (QD) less than 2, strand bias scores (FS) greater than 60, mean mapping quality (MQ) less than 40, or allele mapping bias scores (MQRankSum) less than -12.5 and indels that had QD < 2 or FS > 200. After further filtering variants for biallelic and triallelic markers with genotype calls in at least 50% of individuals, we used GATK ‘VariantsToTable’ to merge SNPs and indels into a single variant table for each cross type.

Finally, we converted our variant tables into AB format, such that the heterozygous parents contribute ‘A’ and ‘B’ alleles to offspring, while the *H. annuus* parent contributes exclusively ‘A’ alleles. At biallelic markers (Fig 2A), sites with two reference alleles became ‘AA’ and sites with the reference allele, and the alternate allele became ‘AB’. At triallelic markers (Fig 2B), sites with the reference allele and one alternate allele became ‘AA’ and sites with the reference allele, and the other alternate allele became ‘AB’. This method randomly assigns ‘A’ and ‘B’ alleles to the homologous chromosomes in each heterozygous parent, so our genetic maps initially consisted of pairs of mirror-imaged linkage groups that we later merged.

### Genetic mapping

We used R/qtl (Broman et al. 2003) in conjunction with R/ASMap (Taylor and Butler 2017) to build genetic maps. After excluding markers with less than 20% or greater than 80% heterozygosity and individuals with less than 50% of markers scored, we used the function ‘mstmap.cross’ with a stringent significance threshold (p.value = 1x10^-16^) to form conservative linkage groups. We used the function ‘plotRF’ to identify pairs of linkage groups with unusually high recombination fractions and the function ‘switchAlleles’ to reverse the genotype scores of one linkage group in each mirrored pair. We did this until reversing genotype scores no longer reduced the number of linkage groups.

Using the corrected genotypes, we made new linkage groups with only the most reliable markers. Namely, we used the function ‘mstmap.cross’ (with the parameter values: dist.fun = “kosambi”, p.value = 1x10^-6^, noMap.size = 2, noMap.dist = 5) on markers with less than 10% missing data and without significant segregation distortion. We refined the resulting linkage groups by removing (1) markers with more than three double crossovers, (2) markers with aberrant segregation patterns (segregation distortion more than two standard deviations above or below the mean segregation distortion of the nearest 20 markers), and (3) linkage groups made up of fewer than four markers.

We progressively pushed markers with increasing amounts of segregation distortion and missing data into the maps using the function ‘pushCross’. After adding each batch of markers, we reordered the linkage groups and dropped markers and linkage groups as described above. Once all the markers had been pushed back, we used the function ‘calc.errorlod’ to identify possible genotyping errors (error scores greater than 2) and replaced those genotypes with missing data. We continued to drop linkage groups, markers, and genotypes that did not meet our criteria until none remained.

Finally, we dropped five excess linkage groups, each made up of fewer than 30 markers, from each map. The markers in these linkage groups mapped to regions of the *H. annuus* genome that were otherwise represented in the final genetic maps but could not be explained by reversed genotypes. Instead, these markers were likely polymorphic in the HA89cms individual used for crosses because of the 2-4% residual heterozygosity in sunflower inbred lines (Mandel et al. 2013).

### SyntR development

To aid in the identification of chromosomal rearrangements, we developed the R package ‘syntR’ (code and documentation available at http://ksamuk.github.io/syntR). This package implements a heuristic algorithm for systematically detecting synteny blocks from marker positions in two genetic maps. The key innovation of the syntR algorithm is coupling a biologically-informed noise reduction method with a cluster identification method better suited for detecting linear (as opposed to circular) clusters of data points.

We based the syntR algorithm on the following statistical and biological properties of genetic maps and chromosomal rearrangements:

1. Synteny blocks appear as contiguous sets of orthologous markers in the same or reversed order in pairs of genetic maps (Pevzner and Tesler 2003, Choi et al. 2007).
2. The inferred order of markers in individual genetic maps is subject to error due to genotyping errors and missing data (Hackett and Broadfoot 2003). This error manifests as slight differences in the order of nearby markers within a linkage group between maps. This mapping error (which we denote ‘error rate one’) results in uncertainty in the sequence of markers in synteny blocks.
3. In genomes with a history of duplication, seemingly orthologous markers can truly represent paralogs. These errors (‘error rate two’) look like tiny translocations and also disrupt marker orders within synteny blocks.
4. When comparing genetic maps derived from genomes without duplications or deletions, every region of each genome will be uniquely represented in the other. Because syntR is made for comparing homoploid genomes with this property, we expect each point in each genetic map to be contained within a single unique synteny block. Therefore, overlaps between synteny blocks are likely errors. Note that this assumption precludes the identification of duplications.
5. Chromosomal rearrangements can be of any size, but smaller rearrangements are difficult to distinguish from error (Pevzner and Tesler 2003). A key decision in synteny block detection is thus the choice of a detection threshold for small rearrangements, which results in a trade-off between error reduction and the minimum size of detectable synteny blocks.

The first step of the syntR algorithm is to smooth over mapping error (error rate one) by identifying highly localized clusters of markers based on a genetic distance threshold (cM) in both maps using hierarchical clustering (Fig 3a). The number of clusters formed is determined by the parameter maximum cluster range (CR_max_) that defines the maximum genetic distance (cM) that any cluster can span in either genetic map. After determining these initial clusters, we smooth the maps by collapsing each multi-marker cluster down into a single representative point (the centroid of the cluster) for processing in subsequent steps. Next, we address errors introduced by poorly mapped or paralogous markers (error rate two) by flagging and removing outlier clusters that do not have a neighboring cluster within a specified maximum genetic distance (cM), a parameter we denote nearest neighbor distance (NN_dist_, Fig 3b).

**Figure 3.**
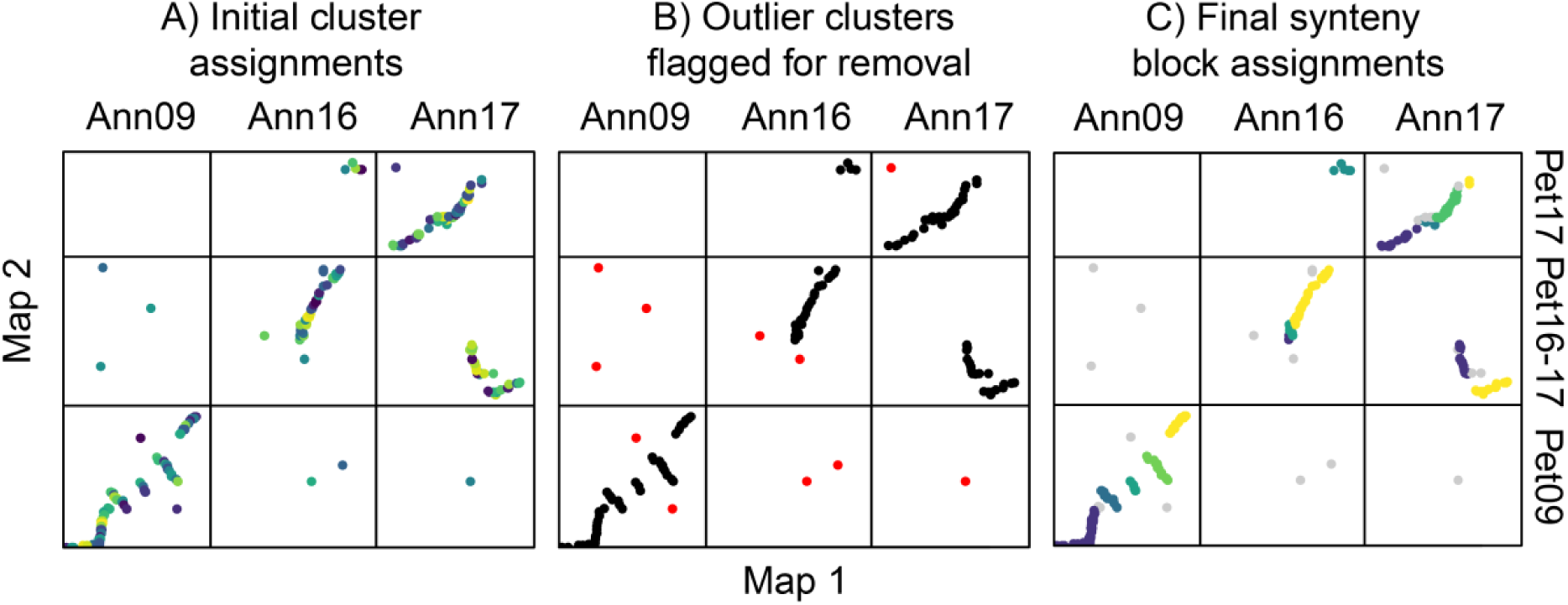
The stages of the syntR algorithm. Each plot shows the relationship between markers or clusters of markers from three chromosomes in two genetic maps. A) Highly localized markers are clustered. Each shade represents an individual cluster of markers that will be collapsed into a single representative point. B) Clusters without another cluster nearby are dropped. Red points represent clusters without a neighbor within 10 cM. C) Clusters are grouped into synteny blocks based on their rank positions. Grey points represent markers that were dropped in previous steps, and each other color represents a different synteny block.

After the noise reduction steps, we define preliminary synteny blocks using a method similar to the “friends-of-friends” clustering algorithm (Huchra and Geller 1982). First, we transform the genetic position of each cluster into rank order to minimize the impact of gaps between markers. We then group clusters that are (1) adjacent in rank position in one of the maps and (2) within two rank positions in the other map (Fig S2). This grouping method further reduces the effect of mapping error by aggregating over pairs (but not triplets) of clusters that have reversed orientations. If a minimum number of clusters per synteny block has been (optionally) defined, we sequentially eliminate blocks that fall below the minimum number of clusters, starting with blocks made up of one cluster and ending with blocks made up of clusters equal to one less than the minimum. After each elimination, we regroup the clusters into new synteny blocks. Finally, we adjust the extents of each synteny block by removing overlapping sections from both synteny blocks so that every position in each genetic map is uniquely represented (Fig 3c).

### Assessing the performance of the syntR algorithm

To evaluate the performance of this method and explore the effect of parameter choice on outcomes, we simulated genetic map comparisons with known inversion breakpoints and error rates in R. The genetic map comparisons were made by randomly placing 200 of markers at 100 positions along a 100 cM chromosome in two maps, reversing marker positions within a defined inversion region in one map, and then repositioning markers based on simulated mapping noise using the following two error parameters: (1) ER1 is the standard deviation of a normal distribution used to pick the distances markers are pushed out of their correct positions (e.g., when ER1 is 1 cM 95% of markers will be within 2 cM of their true position); (2) ER2 is the proportion of markers that are repositioned according to a uniform distribution (i.e., these markers can be moved to any position on the simulated chromosome).

We initially ran syntR using fixed syntR parameters (CR_max_ = 2 and NN_dist_ = 10) on multiple simulated maps, which were made using variable parameters (inversion size: 2.5-50 cM, ER1: 0-2.0 cM, and ER2: 0-20%), and counted the number of times the known breakpoints were identified within 1 cM (Fig S3). As expected, we find that rearrangement size affects the false negative rate (i.e., failing to detect known breakpoints), such that smaller inversions are more likely to be missed (Fig S3c), but does not affect the false positive rate (i.e., detecting breakpoints where there are none). We also find that increasing both types of error in the genetic maps tends to increase both the false positive and false negative rates, although ER1 has a much stronger effect on the false positive rate than any other combination (Fig S3a,b).

Using the same simulation methods as above but now varying the syntR parameter CR_max_, we find that small values of CR_max_ yield high false positive rates while large values yield high false negative rates (Fig S4a). In addition, the ER1 parameter has a strong effect on the relationship between CR_max_ and the false positive rate. Higher values of CR_max_ are needed to reduce the false positive rate when ER1 is also high (Fig S4b). This means that picking an appropriate CR_max_ value is key to the accuracy of this method.

Although NN_dist_ has a much weaker effect on outcomes than CR_max_, it is useful to consider both parameter values carefully.

When the syntR heuristic algorithm is performing well, the final synteny blocks should represent all positions in the two genetic maps being compared (Chen et al. 2009). Based on this characteristic, we developed a method to choose optimal syntR tuning parameters (CR_max_ and NN_dist_) that maximize the representation of the genetic maps and markers in synteny blocks. In this method a user: (1) runs syntR with a range of parameter combinations; (2) saves summary statistics about the genetic distance of each map represented in the synteny blocks and the number of markers retained for each run; and (3) finds the parameter combination that maximizes a composite statistic that equally weights these three measures. In cases where there are multiple local maxima, we suggest choosing the local maximum with the smallest value of CR_max_ to reduce the number of potential false positives.

The “maximize representation” method for choosing syntR parameters has several benefits. First, it does not rely on any additional information (e.g., error rate estimates from the genetic maps compared). Second, when we use this method to choose the best parameters for simulated genetic maps, we find that these parameter values also minimize false positive and false negative rates (Fig S5). Third, when we simulate biologically realistic genetic map comparisons, the absolute value of false positives and false negatives are small. For example, when comparing two genetic maps in which ∼95% of markers are within 1 cM of their true position (ER1 = 0.5) and 5% of markers are randomly permuted (ER2 = 0.05), nonexistent breakpoints will be identified 0.1 times and a breakpoint of a 20 cM inversion will be missed 0.04 times. These low error rates also highlight the overall robustness and accuracy of the syntR algorithm.

In addition to performing simulations, we compared the synteny blocks identified by syntR to those identified by other means in a previously published comparison of *H. niveus* ssp. *tephrodes* and *H. argophyllus* maps to *H. annuus* (Barb et al. 2014). To do this, we formatted the original datasets for input into syntR and used the “maximize representation” method to determine the optimal parameter values for the two comparisons (*H. niveus vs. H. annuus*: CR_max_ = 1.5, NN_dist_ = 30; *H. argophyllus vs. H. annuus*: CR_max_ = 2, NN_dist_ = 20). We found that syntR was in strong agreement with previous work (Fig S6), recovering all the same translocations and most of the same inversions as the Barb et al. (2014) maps. Most of the cases of mismatches were very small or weakly supported inversions in the Barb et al. (2014) maps that syntR did not identify.

### Finding synteny blocks

We used syntR to identify synteny blocks between our newly generated genetic maps and an ultra-high-density map of *H. annuus* that was used to build the sunflower genome that we use as a reference (Badouin et al. 2017). This allowed us to easily convert between physical position in the *H. annuus* reference and position in the *H. annuus* genetic map. Using this property, we further compared two previously published genetic maps for the closely related sunflower species, *H. niveus* ssp. *tephrodes* and *H. argophyllus* (Barb et al. 2014), to the same *H. annuus* map. We aligned marker sequences from the published maps to the *H. annuus* reference using bwa and converted well-aligned markers (MQ > 40) to their positions in the *H. annuus* genetic map.

Initially, we ran syntR using parameters identified through the “maximize representation” method for each map comparison separately (Table S1). However, varying CR_max_ revealed rearrangements that were shared between the maps (Fig S7). Therefore, we ran syntR again using a range of CR_max_ values that included the best fit for each comparison (1.0 - 3.5 in 0.5 increments) and extracted a curated set of synteny blocks from the output. A synteny block was retained if it fulfilled any of the following criteria (in decreasing order of importance): (1) it was found in another species, (2) it was identified in the majority of syntR runs for a single species, (3) it maximized the genetic distance represented by synteny blocks. We present this curated set of synteny blocks below, but our results are unchanged if we use the individually-fit synteny blocks.

**Table 1.**
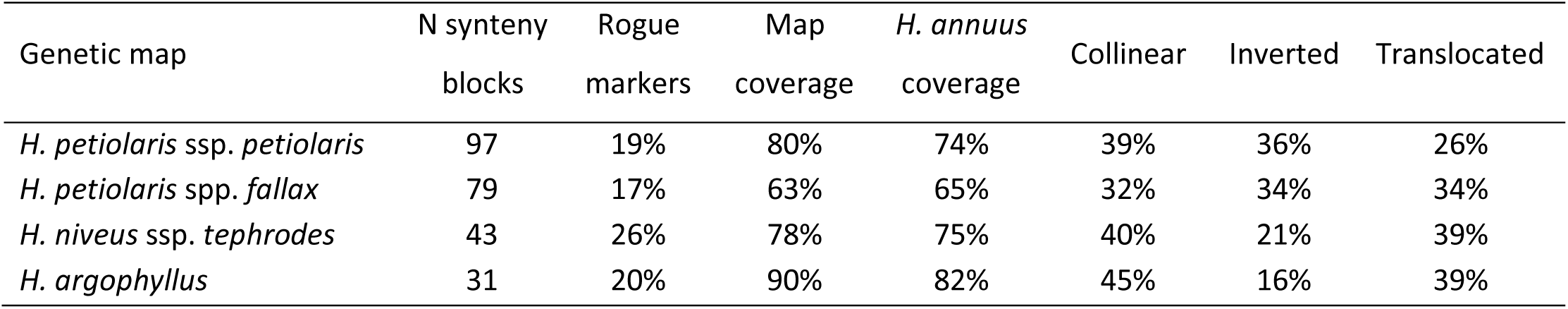
Properties of the synteny blocks found using a syntR analysis between genetic maps of *H. annuus* and four other *Helianthus* taxa. The proportion of rogue markers is based only on the chromosomes without translocations in any map (i.e., chromosomes 1-3, 5, 8-10, 11, and 14). For those chromosomes, the majority of marker mapped to a single *H. annuus* chromosome. The other markers are considered rogue.

| Genetic map | N synteny<br>blocks | Rogue<br>markers | Map<br>coverage | *H. annuus*<br>coverage | Collinear | Inverted | Translocated |
| --- | --- | --- | --- | --- | --- | --- | --- |
| *H. petiolaris* ssp. *petiolaris* | 97 | 19% | 80% | 74% | 39% | 36% | 26% |
| *H. petiolaris* spp. *fallax* | 79 | 17% | 63% | 65% | 32% | 34% | 34% |
| *H. niveus* ssp. *tephrodes* | 43 | 26% | 78% | 75% | 40% | 21% | 39% |
| *H. argophyllus* | 31 | 20% | 90% | 82% | 45% | 16% | 39% |

We named the chromosomes in our genetic maps based on their synteny with the standard order and orientation of *H. annuus* chromosomes (Tang et al. 2002, Bowers et al. 2012) following Barb et al. (2014) but with shortened prefixes (A = *H. annuus*, R = *H. argophyllus*, N = *H. niveus* ssp. *tephrodes*, P = *H. petiolaris* ssp. *petiolaris*, F = *H. petiolaris* ssp. *fallax*). For example, an *H. petiolaris* ssp. *fallax* chromosome made up of regions that are syntenic with *H. annuus* chromosomes 4 and 7 is called F4-7.

### Karyotype reconstruction and analysis

We used our inferred synteny blocks and the software MGR v 2.01 (Bourque and Pevzner 2002) to infer ancestral karyotypes for our five *Helianthus* taxa and to determine the number of chromosomal rearrangements that occurred along each branch of the species tree. To run the MGR analysis, we needed the order and orientations of synteny blocks in all five maps. However, individual synteny blocks were often missing from one or more of our final maps. We approached this problem in two ways. First, we inferred the likely position of missing synteny blocks based on the location of markers that were too sparse to be grouped by syntR and matched the location of synteny blocks in other maps. In the second case, we dropped any synteny blocks that were not universally represented.

Because we already had two sets of synteny blocks for each map (curated and individually optimized), we ran the MGR analyses using three different sets of synteny blocks: (set 1) curated and inferred, (set 2) curated and present in all five maps, (set 3) individually optimized and present in all five maps.

### Data availability

The R program, syntR, is available on GitHub: https://github.com/ksamuk/syntR. The sequences used to generate genetic maps are available on the SRA: http://www.ncbi.nlm.nih.gov/bioproject/598366. All other data and scripts are available on dryad: https://doi.org/10.5061/dryad.7sqv9s4pc.

## Results

### Genetic maps

Both *H. petiolaris* genetic maps are made up of the expected 17 chromosomes and have very high marker density (Fig 4, Fig S8). Only 6% of the *H. petiolaris* ssp. *petiolaris* map and 10% of the *H. petiolaris* ssp. *fallax* map fails to have a marker within 2 cM (Fig S9). Overall, both maps are somewhat longer than the *H. petiolaris* map reported by Burke et al. (2004). Although this could represent real variation between genotypes, it could also be the result of spurious crossovers that are inferred based on genotyping errors. Because genotyping errors are proportional to the number of markers, maps with high marker densities are more likely to be inflated. Indeed, building maps with variants that were thinned to 1 per 150 bp using vcftools version 0.1.13 (Danecek et al. 2011) yields collinear maps that are closer to the expected lengths (Table S2, Fig S10). We present subsequent results based on the full maps to improve our resolution for detecting small rearrangements.

**Figure 4.**
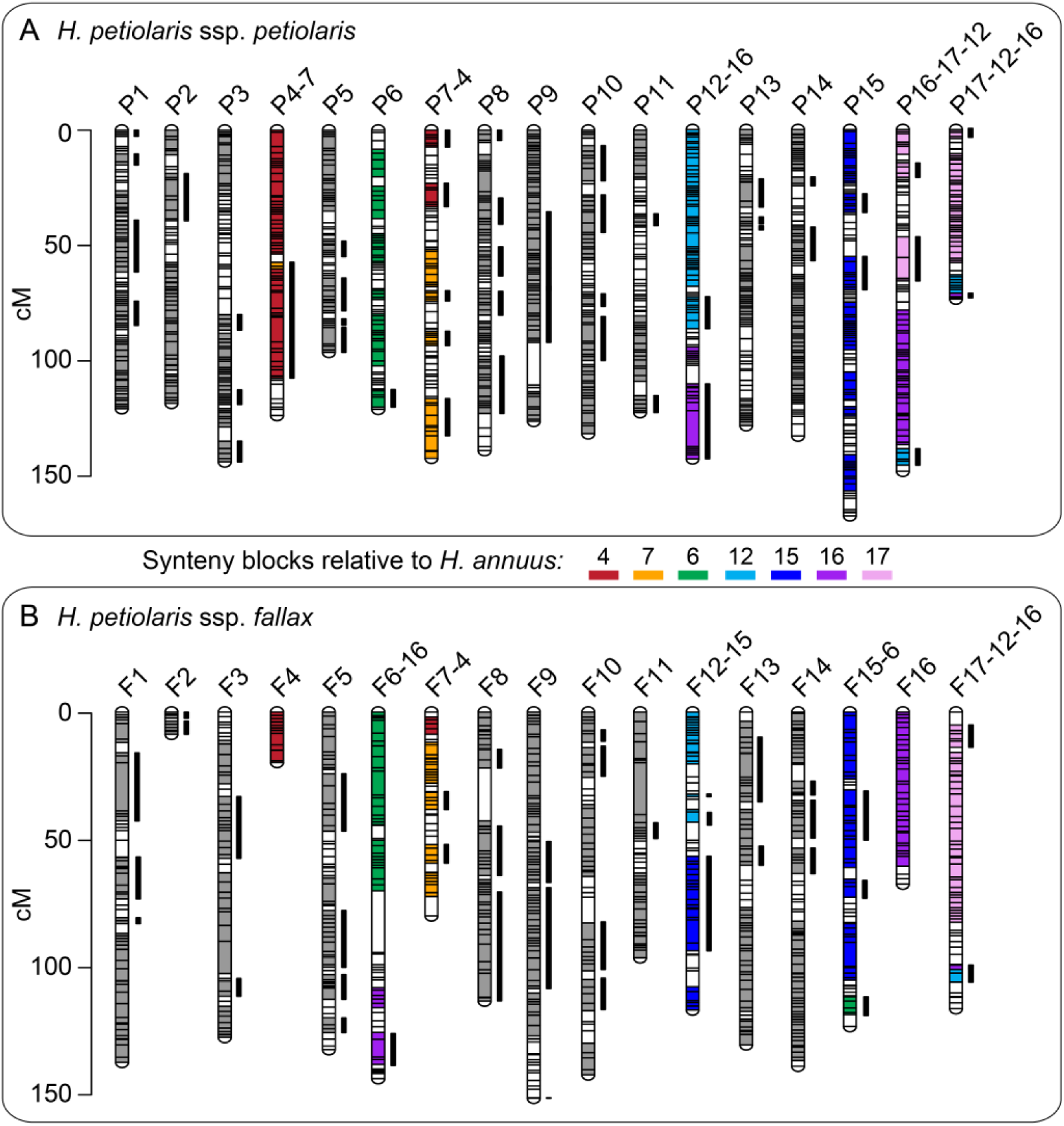
*Helianthus petiolaris* genetic maps showing blocks of synteny with *H. annuus*. Each horizontal bar represents a genetic marker. The thick vertical bars next to chromosomes represent synteny blocks that are inverted relative to the *H. annuus* genetic map. Where there are no translocations between *H. petiolaris* and *H. annuus* chromosomes (e.g.. all synteny blocks in P1 and F1 are syntenic with A1), the synteny blocks are shown in grey. Where there are translocations, the synteny blocks are color-coded based on their synteny with *H. annuus* chromosomes. Regions that are not assigned to a synteny block remain white. The synteny blocks plotted are those curated based on multiple runs of syntR using different parameters. Please see Fig S12 for a labeled version. This figure was made with LinkageMapView (Ouellette et al. 2017).

Despite the general expansion of our maps, we find that chromosomes 2 and 4 in the *H. petiolaris* ssp. *fallax* map (F2 and F4) are unexpectedly short (Fig 4). When we look at the distribution of markers for this map relative to the *H. annuus* reference, we find very few variable sites in the distal half of these chromosomes (Fig S11). That is, this individual was homozygous along vast stretches of F2 and F4.

These runs of homozygosity could be explained by recent common ancestry (i.e., inbreeding) or a lack of variation in the population (e.g, because of background selection or a recent selective sweep).

Regardless, the lack of variable sites within the *H. petiolaris* ssp. *fallax* individual used for crosses explains the shortness of F2 and F4. Notably, we find the same pattern on the distal half of *H. annuus* chromosome 7 and find that this region is also not represented in the *H. petiolaris* spp. *fallax* map.

### Synteny blocks

Using syntR, we recovered 97 genetic regions that are syntenic between the *H. petiolaris* ssp. *petiolaris* and *H. annuus* and 79 genetic regions that are syntenic between the *H. petiolaris* ssp. *fallax* and *H. annuus* (Fig 4). We also recovered synteny blocks for the *H. niveus* ssp. *tephrodes* and *H. argophyllus* comparisons that are similar to those found previously (Fig S13). In all four comparisons, syntR successfully identified synteny blocks that cover large proportions (63%-90%) of each genetic map even in the face of a very high proportion of markers that map to a different chromosome than their neighbors (Table 1). These “rogue markers” could be the result of very small translocations, poorly mapped markers, or extensive paralogy. Over and above the prevalence of rogue markers, the karyotypes we recovered are substantially rearranged. Only between 32% and 45% of synteny blocks for each map are collinear with the *H. annuus* genetic map in direct comparisons (Table 1).

### Karyotype reconstruction and chromosomal rearrangement

Because nested and shared rearrangements can obscure patterns of chromosome evolution, we use the MGR analyses to predict the most likely sequence of rearrangements in a phylogenetic context before quantifying the rearrangement rate. These MGR analyses identified similar patterns of chromosome evolution regardless of the exact set of synteny blocks that we used (Table S5). Multiple taxa share many rearrangements, and the similarity of karyotypes matches known phylogenetic relationships. Moreover, MGR analyses run without a guide tree inferred the known species tree, and MGR analyses run with all other topologies identified an inflated number of chromosomal rearrangements.

Using the most complete set of synteny blocks (set 1), we find that 88 chromosomal rearrangements occurred across the phylogeny (Fig 5). Then, using the most current divergence time estimates for this group (Todesco et al. 2019) and conservatively assuming that *H. niveus* ssp. *tephrodes* diverged at the earliest possible point, we estimate that 7.9 (7.8-8) rearrangements occurred per million years in this clade (Tables S3-S5). To further explore the potential range of rearrangement rates, we considered other estimates of divergence times in sunflower (Sambatti et al. 2012, Mason 2018) and the other sets of synteny blocks. Overall, the lowest rate we identified was 2.6 rearrangements per million years, while the highest rate was indeterminable because some minimum divergence time estimates for the group include 0 (Tables S3-S5).

**Figure 5.**
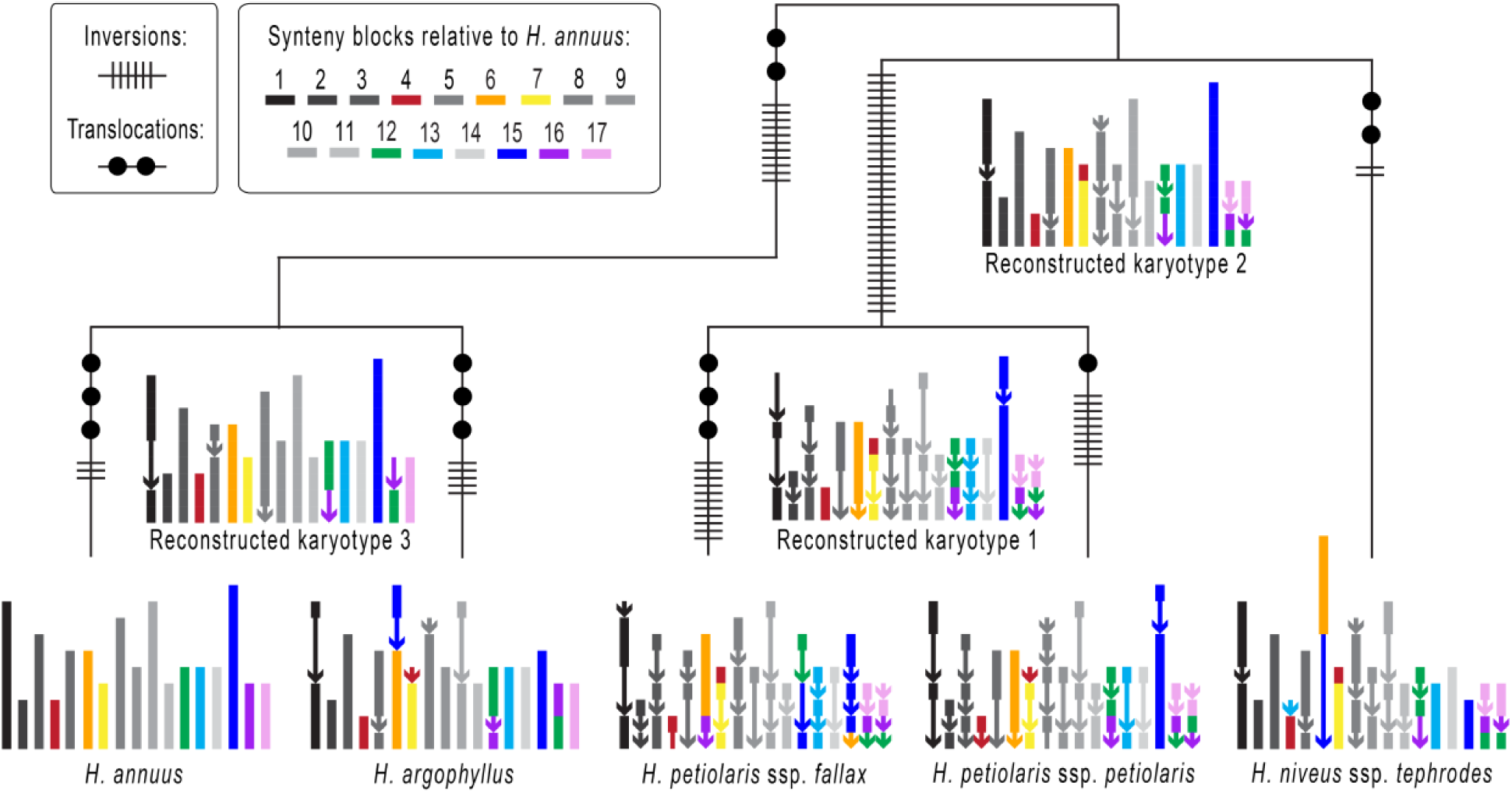
Diagram showing the karyotypes of 5 *Helianthus* taxa as well as reconstructed ancestral karyotypes and the locations of chromosomal rearrangements. The karyotypes were built using synteny block set 1, which were curated based on multiple syntR runs and inferred when missing. Each synteny block is represented using a line segment that is color-coded based on its position in the *H. annuus* genome (see Fig S14 for a labeled version). Chromosomes without translocations in any map are plotted in grey, and synteny blocks that are inverted relative to *H. annuus* are plotted using arrows. Also, note that along some branches the same pair of chromosomes is involved in multiple translocations.

The 88 rearrangements include 74 inversions and 14 translocations that are quite evenly distributed across the phylogeny. However, the excess inversions indicate that it is unlikely that the rate of inversions is equal to the rate of translocation (binomial test, 5.1x10^-11^). Furthermore, we find that only 8 of the 17 chromosomes are involved in the 14 translocations we identified. If translocations were equally likely for all chromosomes, this asymmetry is very unlikely to have happened by chance (the probability of sampling ≤ 8 chromosomes in 14 translocations is 8.0x10^-8^, Fig S15), suggesting that some chromosomes are more likely to be involved in translocations than other. In line with this observation, we see that some chromosome segments are repeatedly translocated. For example, A4 and A7 are involved in several exchanges, and part of A6 has a different position in almost every map (Fig 5).

## Discussion

Large-scale chromosomal changes may be key contributors to the process of adaptation and speciation, yet we still have a poor understanding of rates of chromosomal rearrangement and the evolutionary forces underlying those rates. Here, we devised a novel, systematic method for comparing any pair of genetic maps, and performed a comprehensive analysis of the evolution of chromosomal rearrangements in a clade of sunflowers. We created two new genetic maps for *Helianthus* species and used our new method to identify a wide range of karyotypic variation in our new maps, as well as previously published maps. Consistent with previous studies, we discovered a high rate of chromosomal evolution in the annual sunflowers. Further, we found that inversions are more common than translocations and that certain chromosomes are more likely to be translocated. Below, we discuss the evolutionary and methodological implications of this work and suggest some next steps in understanding the dynamic process of chromosomal rearrangement.

### Identifying rearrangements

Studying the evolution of chromosomal rearrangements requires dense genetic maps and systematic methods to analyze and compare these maps between species. Our new software, syntR, provides an end-to-end solution for systematic and repeatable identification of synteny blocks in pairs of genetic maps with any marker density. Our tests on real and simulated data find that syntR recovers chromosomal rearrangements identified previous by both manual comparisons and cytological study, suggesting that syntR is providing an accurate view of karyotypic differences between species.

Overall, we believe syntR will be a valuable tool for the systematic study of chromosomal rearrangements in any species. The only data syntR needs to identify synteny blocks is relative marker positions in two genetic maps. This fact is significant because, although the number of species with whole genome sequence and methods to detect synteny blocks from those sequences are rapidly accumulating, such as Mauve (Darling et al. 2004), Cinteny (Sinha and Meller 2007), syMAP (Soderlund et al. 2011), SynChro (Drillion et al. 2014) and SyRI (Goel et al. 2019), it is still uncommon to have multiple closely related whole genome sequences that are of sufficient quality to compare for karyotype differences. At the same time, the proliferation of reduced representation genome sequencing methods means that it is easy to generate many genetic markers for non-model species and produce very dense genetic maps. Furthermore, syntR allows comparisons to include older genetic map data that would otherwise go unused. The simplicity of the syntR algorithm will facilitate rapid karyotype mapping in a wide range of taxa.

We also believe that syntR provides a baseline for the development of further computational and statistical methods for the study of chromosomal rearrangements. One fruitful direction would be to integrate the syntR algorithm for synteny block detection directly into the genetic map building process (much like GOOGA, Flagel et al. 2019). Another key extension would be to allow syntR to compare multiple genetic maps simultaneously to detect synteny blocks in a group of species (e.g., by leveraging information across species). Finally, formal statistical methods for evaluating the model fit and the uncertainty involved with any set of synteny blocks would be a major (albeit challenging) improvement to all existing methods, including syntR.

### The similarity of *H. petiolaris* maps to previous studies

Compared with previous work, we found more inversions and fewer translocations between *H. petiolaris* subspecies and *H. annuus* (Rieseberg et al. 1995, Burke et al. 2004). This is probably due to a combination of factors. First, there appears to be karyotypic variation within some *Helianthus* species (Heiser 1948, Heiser 1961, Chandler et al. 1986). Second, the maps presented here are made up of more markers and individuals, which allowed us to identify small inversions that were previously undetected as well as to eliminate false linkages that can be problematic in small mapping populations. Lastly, we required more evidence to call rearrangements. Although we recovered some of the translocations supported by multiple markers in Rieseberg et al. (1995) and Burke et al. (2004), we did not recover any of the translocations supported by only a single sequence-based marker. Given the high proportion of “rogue markers” in our maps, it is likely that some of the putative translocations recovered in those earlier comparisons are the result of the same phenomenon.

On the other hand, we found that rearrangements between our *H. petiolaris* maps match the translocations predicted from cytological studies quite well. Heiser (1961) predicted that *H. petiolaris* ssp. *petiolaris* and *H. petiolaris* ssp. *fallax* karyotypes would have three chromosomes involved in two translocations that form a ring during pairing at meiosis, as well as the possibility of a second independent rearrangement. This exact configuration is likely to occur at meiosis in hybrids between the *H. petiolaris* subspecies maps we present here (Fig S16). Also, the most noteworthy chromosome configuration in cytological studies of *H. annuus*-*H. petiolaris* hybrids (Heiser 1947, Whelan 1979, Ferriera 1980, Chandler et al. 1986) was a hexavalent (a six-chromosome structure) plus a quadrivalent (a four-chromosome structure). Again, this is the configuration that we would expect in a hybrid between *H. annuus* and the *H. petiolaris* ssp. *petiolaris* individual mapped here. Furthermore, the complicated arrangement and relatively small size of A12, A16 and A17 synteny blocks in *H. petiolaris* might explain why cytological configurations in *H. annuus*-*H. petiolaris* hybrids are so variable.

Interestingly, the rearrangements identified between *H. argophyllus* and *H. annuus* karyotypes here and in Barb et al. (2014) also match the cytological studies better than an earlier comparison of sparse genetic maps (Heesacker et al. 2009). It seems that, in systems with the potential for high proportions of rogue markers, many markers are needed to identify chromosomal rearrangements reliably.

### Total rearrangement rates

Our data suggest that annual sunflowers experience approximately 7.9 chromosomal rearrangements per million years. This rate overlaps with recent estimates for this group (7.4-10.3, Barb et al. 2014) and is even higher than the estimate that highlighted sunflower as a group with exceptionally fast chromosomal evolution (5.5-7.3, Burke et al. 2004). However, since Burke et al. (2004), chromosomal rearrangements have been tracked in many additional groups, including mammals (Ferguson-Smith and Trifonov 2007, Martinez et al. 2016, da Silva et al. 2019), fish (Molina et al. 2014, Ayres-Alves et al. 2017), insects (Rueppell et al. 2016, Corbett-Detig et al. 2019), fungi (Sun et al. 2017) and plants (Yogeeswaran et al. 2005, Schranz et al. 2006, Huang et al. 2009, Vogel et al. 2010, Latta et al. 2019). Of these analyses, relatively few have systematically studied karyotypes evolution across multiple species and estimated total rearrangement rates. Of those that do, most studies report less than 7.9 chromosomal rearrangements per million years, for example, in *Solanum* (0.36-1.44, Wu and Tanksley 2010), *Drosophila* (0.44-2.74, Bhutkar et al. 2008) and mammals (0.05-2.76, Murphy et al. 2005). But there are exceptions, such as a comparison of genome sequences that revealed up to 35.7 rearrangements per million years in some grass lineages (Dvorak et al. 2018).

At the same time, we are likely underestimating rearrangement rates here for two reasons. First, we used conservative thresholds for calling rearrangements. For example, some proportion of the rogue markers that we identified could be the result of very small but real chromosomal rearrangements. Second, our ability to resolve very small synteny blocks and breakpoints between synteny blocks depends on marker density. Until we have full genome sequences to compare (like for the grass lineages), we could be failing to detect very small rearrangements and falsely inferring that independent rearrangements are shared. However, regardless of just how much we are underestimating the rate, sunflower chromosomes are evolving quickly. This high rate of chromosomal evolution could be a consequence of a higher rate of chromosomal mutation, a decreased chance that chromosomal polymorphisms are lost, or both processes.

### Type of rearrangements

We found that inversions and interchromosomal translocations dominate chromosomal evolution in *Helianthus.* This pattern is common in angiosperm lineages (Weiss-Schneeweiss and Schneeweis 2012) and fits with the consistent chromosome counts across annual sunflowers (2n = 34, Chandler et al. 1986). In addition, we found more inversions than translocations, which has previously been seen in both plant (Wu and Tanksley 2010, Amores et al. 2014) and animal systems (Rueppell et al. 2016) and echoes general reports that intrachromosomal rearrangements are more common than interchromosomal rearrangements (Pevzner and Tesler 2003). These consistent rate differences are notable because, although both rearrangement types depend on double strand breaks, two of the major consequences of chromosomal rearrangements, underdominance (i.e., rearrangement heterozygotes are less fit than either homozygote) and recombination modification, might be more common for some types of rearrangements.

Translocations have a more predictable effect on hybrid fertility, while inversions consistently reduce recombination. Reciprocal translocation heterozygotes can affect fertility because missegregation during meiosis can cause half of the gametes to be unbalanced and thus inviable (White 1973, King 1993). Although inversion heterozygotes can also produce unbalanced gametes, whether that happens is dependent on the size of the inversion and whether disrupted pairing during meiosis inhibits crossovers (Searle 1993). When inversions are small or have suppressed crossing over, they will not be strongly underdominant. On the other hand, inversions often exhibit reduced recombination either because recombination is suppressed through disrupted pairing (Searle 1993) or ineffective through the production of inviable gametes (Rieseberg 2001). While interactions between reduced recombination and adaptation with gene flow have been extensively examined in the case of inversions (Kirkpatrick and Barton 2006, Hoffman and Rieseberg 2008, Yeaman and Whitlock 2011, Yeaman 2013), it is not clear whether the same pattern will be common for translocations (but see Fishman et al. 2013, Stathos and Fishman 2014 for one example). Translocations bring together previously unlinked alleles and mispairing at translocation breakpoints could suppress crossing over, but recombination inside reciprocal translocations will not necessarily produce inviable gametes and thus reduce effective recombination.

Although any selective force could be responsible for the evolution of any chromosomal rearrangement, potential differences in the relative magnitude of underdominance versus recombination suppression may contribute to the evolution of sunflower chromosomes. While many chromosomal rearrangements in sunflowers appear to be strongly underdominant (Chandler 1986, Lai et al. 2005), inversions typically are not (L. Rieseberg, unpublished). If translocations tend to be more underdominant than inversions, they would be less likely to evolve through drift and more likely to cause reproductive isolation directly. This could explain why translocations are less common than inversions and why pollen viability is accurately predicted by the number of translocations inferred from cytological studies (Chandler et al. 1986). At the same time, recent genomic analyses have identified several extensive regions of very low recombination caused by large inversions segregating in natural sunflower populations (Todesco et al. 2019, Huang et al. 2019). Mutations that segregate for extended periods are unlikely to be strongly underdominant, and these inversions are associated with multiple adaptive alleles (Todesco et al. 2019), which is consistent with a role for selection in their origin or maintenance.

### Non-random chromosomal rearrangement

We also found that some sunflower chromosomes are involved in more translocations than others. This pattern has been observed in wheat (Badaeva et al. 2007) and breakpoint reuse is a common phenomenon in comparative studies of karyotypes (Pevzner and Tesler 2003, Bailey et al. 2004, Murphy et al. 2005, Larkin et al. 2009). Many studies support the idea that chromosomal regions with greater sequence similarity are more likely to recombine and thus potentially generate novel chromosomal arrangements. Some of the clearest examples of this come from the polyploidy literature, where chromosomes with ancestral homology are more likely to recombine (Nicolas et al. 2007, Marone et al. 2012, Mason et al. 2014, Tennessen et al. 2014, Nguepjop et al. 2016). However, centromeres and other repetitive regions can also affect the rate of mutations that cause chromosomal rearrangements (Hardison et al. 2003, Murphy et al. 2005, Raskina et al. 2008, Molnár et al. 2010, Vitte et al. 2014, Ayers-Alves et al. 2017, Li et al. 2017, Corbett-Detig et al. 2019). Given that sunflowers have several genome duplications and a burst of transposable element activity in their evolutionary history (Barker et al. 2008, Kawakami et al. 2011, Staton et al. 2012, Barker et al. 2016, Badouin et al. 2017) it is plausible that ancestral homology or repeat content could be associated with translocation propensity.

Of the above possibilities, an association between repeated translocations and centromeres would be particularly compelling. Beyond the repeat content of centromeres explaining non-random mutation (Kawabe et al. 2006, Sun et al. 2017, but see Lin et al. 2018, Okita et al. 2019), the position and size of centromeres on chromosomes is known to affect meiotic drive and thus the repositioning of centromeres through rearrangement could cause non-random fixation of translocations (Kaszás et al. 1998, Chmátal et al. 2014, Zanders et al. 2014). The relative placement of centromeres has been associated with chromosome evolution in *Brassica* (Schranz et al. 2006) and wheat (Badaeva et al. 2007), and associations between meiotic drive and chromosome evolution have been found in several animal taxa (Bidau and Martí 2004, Palestis et al. 2004, Molina et al. 2014, Blackmon et al. 2019). In sunflower, we see some hints that centromeric repeats might be associated with repeated translocation. Using the locations of the centromere-specific retrotransposon sequence, HaCEN-LINE (Nagaki et al. 2015), to roughly identify the locations of centromeres in our reference, we find that some rearrangement breakpoints, for example, the section of A16 with a different position in each map, are close to putative centromeres (Fig S17-S20). Although a more thorough analysis of centromeric repeat locations and their association with rearrangement breakpoints is required to draw firm conclusions about the importance of centromeres to chromosomal evolution in sunflower, the development of reference sequences for wild sunflower species is underway, which will allow those and other associations to be confirmed. Further, it is time to directly test for meiotic drive in this system by examining the transmission of rearrangements that affect centromeres in gametes produced by plants that have heterozygous karyotypes.

## Conclusion

Understanding the evolution of chromosomal rearrangements remains a key challenge in evolutionary genetics. By developing new software to systematically detect synteny blocks and building new genetic maps, we show that sunflowers exhibit rapid and non-random patterns of chromosomal evolution.

These data generate specific and testable hypotheses about chromosomal evolution in sunflower. We believe that our work will spur additional studies of karyotypic evolution and diversity, and ultimately lead to a more comprehensive understanding of the interplay between chromosomal evolution and speciation.

## Acknowledgments

We thank Jessica Barb for providing marker sequence data, Marcy Uyenoyama for help with our random walk analysis, Greg Baute for sharing hybrid seed, Chris Grassa for growing seedlings and sharing scripts, and both Marco Todesco and Nadia Chaidir for help in the lab. We also thank Jenn Coughlan, Andrew MacDonald, Brook Moyers, Mariano Alvarez, Dolph Schluter, Darren Irwin, Sally Otto, and three anonymous reviewers for thoughtful discussions and help with earlier drafts of this manuscript. This work was supported by an NSERC Postgraduate Scholarship awarded to KLO and an NSERC Discovery Grant awarded to LHR (327475).

## Author contributions

KLO and LHR planned the study. KLO and KS designed and built the R package syntR. KLO made genetic maps, carried out data analysis, and drafted the manuscript. All authors read, edited, and approved the final manuscript.

