## Supplemental material for "Ancestral reconstruction of sunflower karyotypes reveals non-random chromosomal evolution"

This PDF file contains the following supplementary materials:

Figures:
Fig S1: Range Map ........ 2
Fig S2: SyntR clustering diagram ........ 3
Fig S3: SyntR performance ........ 3
Fig S4: Effect of $CR_{max}$ on syntR performance ........ 4
Fig S5: "Maximize representation" performance ........ 4
Fig S6: Synteny blocks from syntR match blocks from previous studies ........ 5
Fig S7: Synteny blocks found using variable parameters ........ 6
Fig S8: Genetic maps with marker density ........ 7
Fig S9: Cumulative marker density plots ........ 8
Fig S10: Full versus thin maps ........ 9
Fig S11: *H. petiolaris* ssp. *fallax* markers ........ 10
Fig S12: *H. petiolaris* maps with synteny blocks ........ 11
Fig S13: *H. niveus* and *H. agrophyllus* maps with synteny blocks ........ 12
Fig S14: Reconstruction of ancestral *Helianthus* karyotypes ........ 13
Fig S15: Non-random translocation probabilities distributions ........ 16
Fig S16: Chromosome pairing at meiosis ........ 17
Fig S17: *H. petiolaris* ssp. *petiolaris* dot plots with centromeric sequence ........ 18
Fig S18: *H. petiolaris* ssp. *fallax* dot plots with centromeric sequence ........ 19
Fig S19: *H. niveus* ssp. *tephrodes* dot plots with centromeric sequence ........ 20
Fig S20: *H. argophyllus* dot plots with centromeric sequence locations ........ 21

Tables:

#### Figure S1 – Range Map

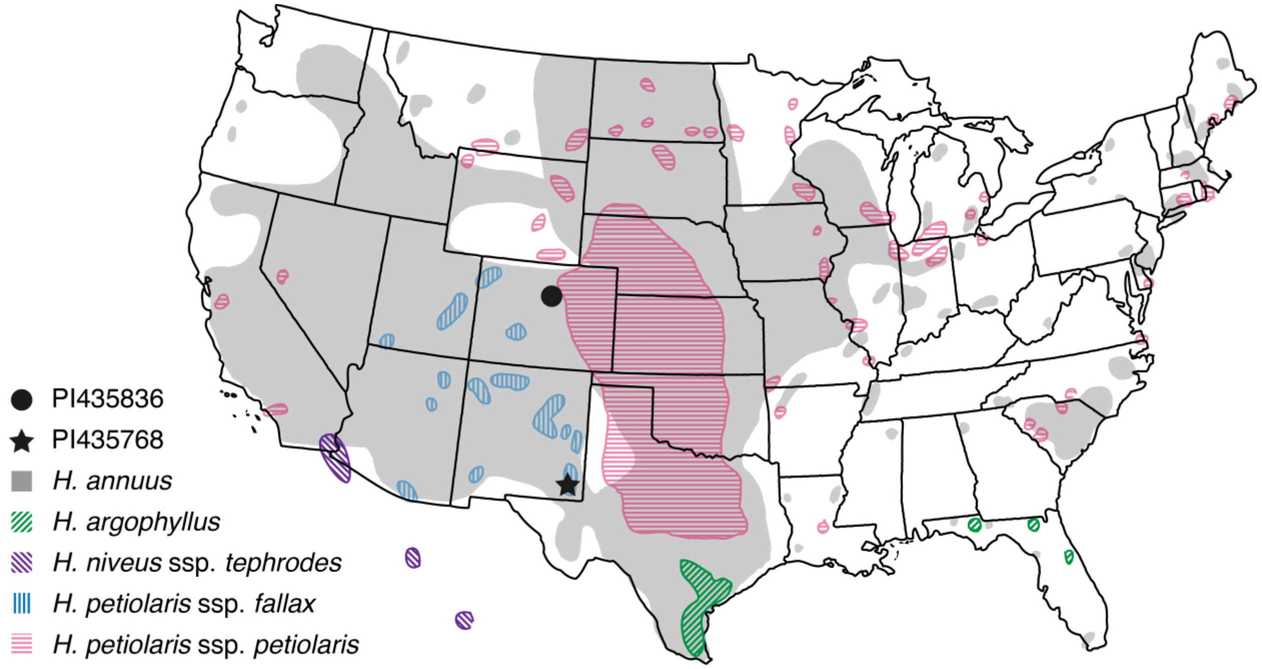

Figure S1 – Approximate range distributions and collection locations for the *Helianthus* taxa used in this study. Redrawn from Rogers et al. 1982.

#### Figure S2 – SyntR clustering diagram

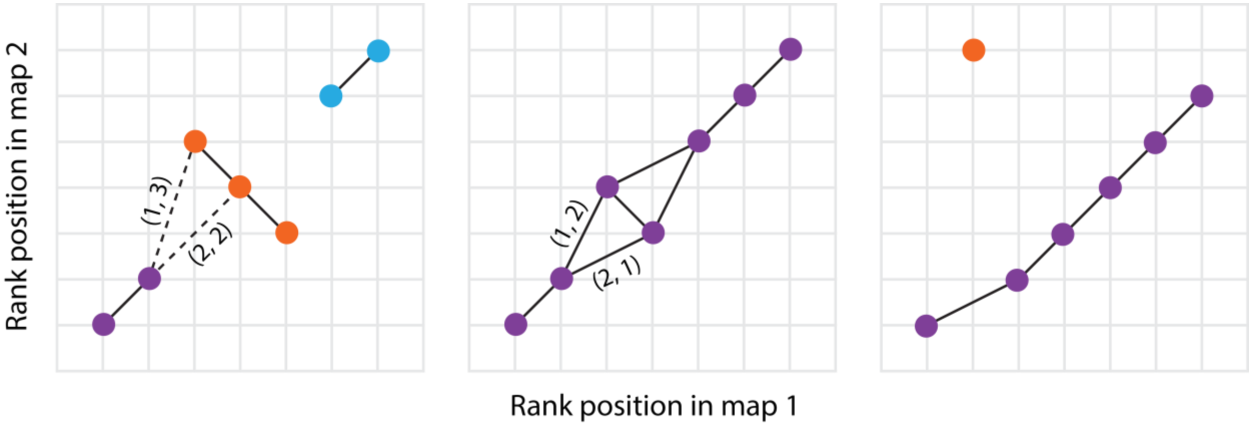

Figure S2 – Illustration of the way syntR groups clusters of markers into synteny blocks. Clusters are grouped if they lie within one rank in one map and within two ranks in the other map. Clusters that fit this criterion are connected by solid lines, while examples of clusters that do not fit this criterion are connected by dashed lines. Numbers in parentheses note the rank distance between points in each genetic map (rank distance in map 1 is given first). Each group of clusters that would make up a synteny block has a unique color.

#### Figure S3 – SyntR performance

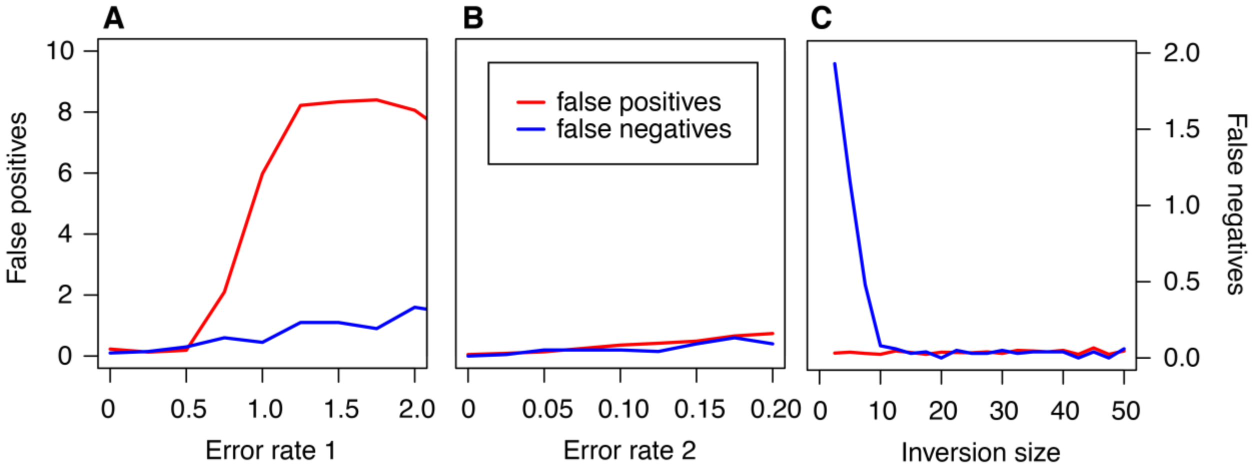

Figure S3 - The number of times simulated breakpoints are incorrectly identified (false positives) or missed (false negatives) by syntR. Each position represents the mean number of errors after running syntR on 100 simulated map comparisons, where breakpoints are considered correctly identified if they were within 1 cM of a known breakpoint. For each run, the syntR parameters were fixed ($CR_{max}$ = 2 and $NN_{dist}$ = 10) and two of three map parameters were fixed ($ER_1$ = 0.5, $ER_2$ = 0.1, inversion size = 20 cM) while the third was varied ($0 \leq ER_1 \leq 2.0$, $0 \leq ER_2 \leq 0.2$, 2.5 cM $\leq$ inversion size $\leq$ 50 cM).

#### Figure S4 – Effect of $CR_{max}$ syntR performance

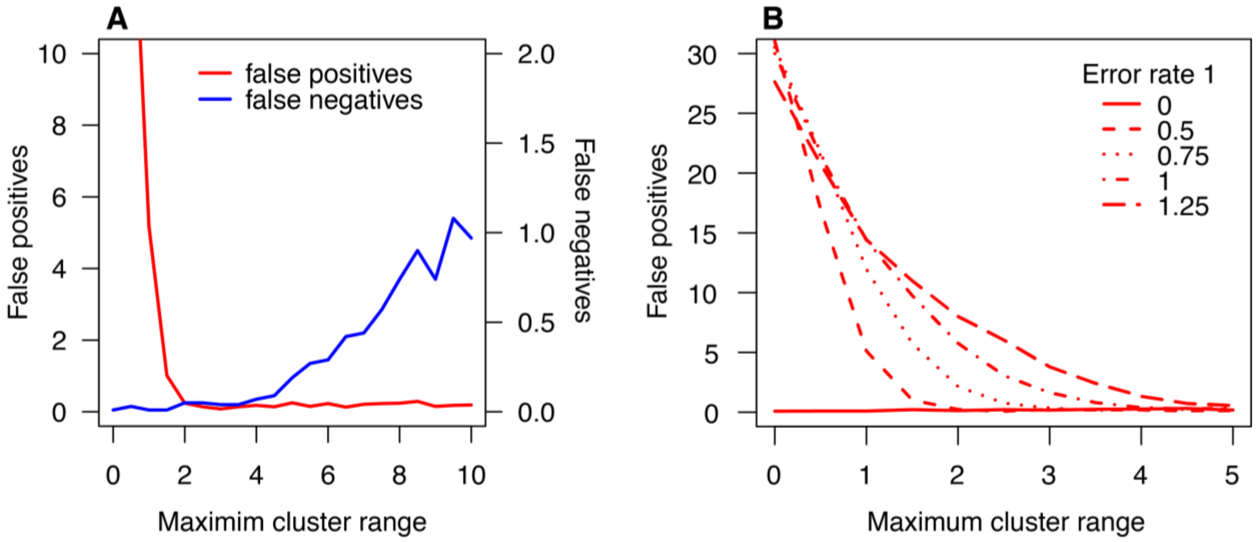

Figure S4 – The effect of $CR_{max}$ on the number of times simulated breakpoints are incorrectly identified (false positives) or missed (false negatives) by syntR. Each point represents the mean number of errors after running syntR on 100 simulated map comparisons, where breakpoints are considered correctly identified if they were within 1 cM of a known breakpoint. For each run, the following parameters were fixed: $NN_{dist}$ = 10, $ER_1$ = 0.5 (only in A), $ER_2$ = 0.1, inversion size = 20 cM.

#### Figure S5 – "Maximize representation" performance

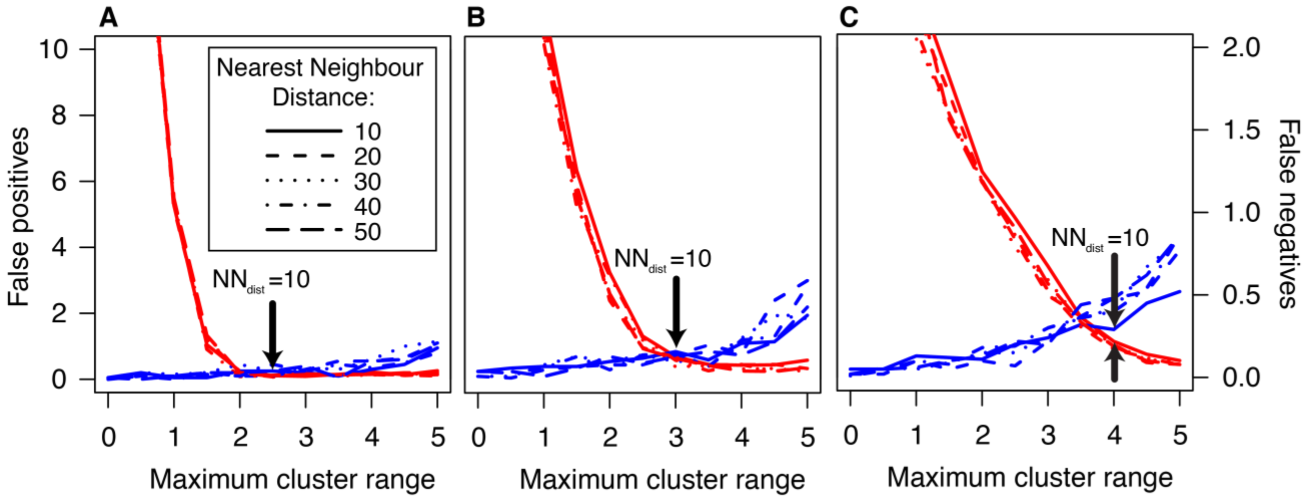

Figure S5 – The number of times simulated breakpoints are incorrectly identified (false positives) or missed (false negatives) by syntR in comparisons with increasing amounts of error in the simulated maps. Arrows point to the syntR parameter values chosen by the maximize representation method. Each panel representes simulated maps with a 20 cM inversion and the following error parameters: A) $ER_1$ = 0.5, $ER_2$ = 0.05; B) $ER_1$ = 0.8, $ER_2$ = 0.1; C) $ER_1$ = 1, $ER_2$ = 0.1.

#### Figure S6 – Synteny blocks from syntR match blocks from previous studies

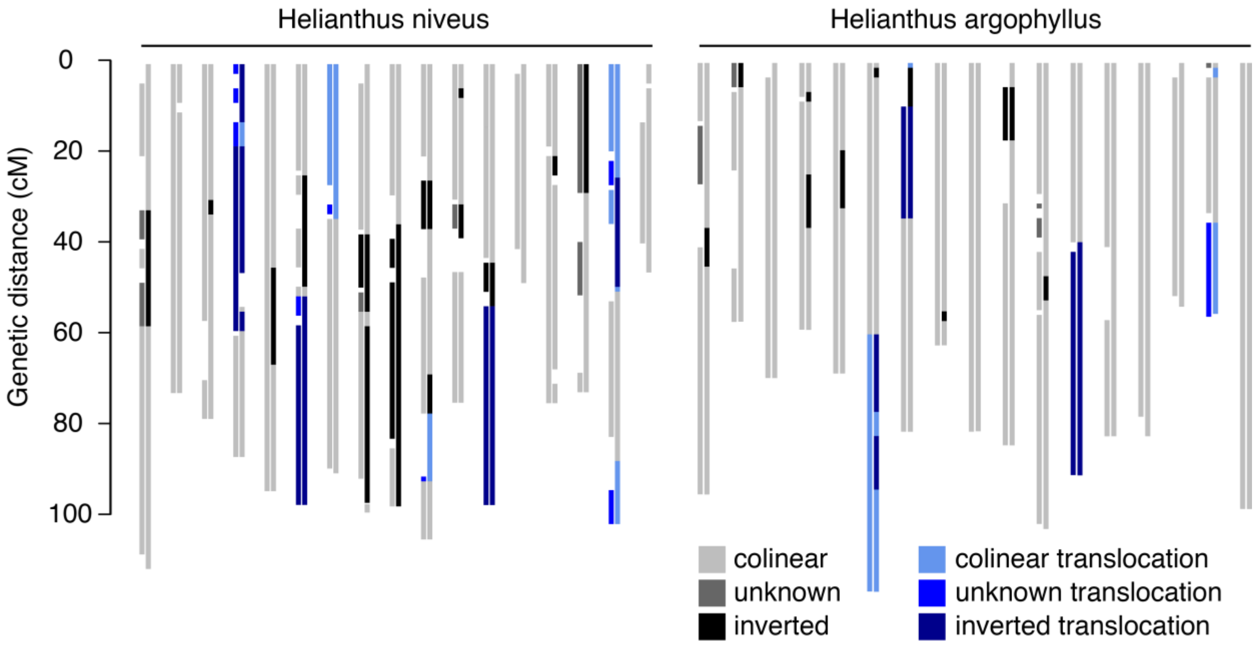

Figure S6 - Match between synteny blocks detection and categorization using different methods. Each pair of chromosomes shows the synteny blocks found using syntR (left) and identified in Barb et al. 2014 (right). The synteny blocks are colored based on whether they are translocated and whether they are inverted or have an unknown orientation relative to *H. annuus*. The amount of genetic distance for which both methods produce matching categorizations are as follows: *H. niveus*: 75% matching (7% partial, where partial matches are those between colinaer/inverted and unknown segments or colinear/inverted translocations and unknown translocations), 17% missing in one map, and 8% mismatched; *H. argophyllus*: 86% matching (3% partial), 9% missing, and 5% mismatched.

#### Table S1 – SyntR parameters used

Table S1 – The syntR tuning parameters used to identify synteny blocks between *H. annuus* and several other genetic maps.

| Taxon | $CR_{max}$ | $NN_{dist}$ | min_block_size |
|---|---|---|---|
| *H. petiolaris* ssp. *petiolaris* | 3.0 | 10 | 3 |
| *H. petiolaris* ssp. *fallax* | 2.5 | 10 | 3 |
| *H. niveus* ssp. *tephrodes* | 1.5 | 10 | 3 |
| *H. argophyllus* | 1.5 | 10 | 3 |

#### Figure S7 – Synteny blocks found using variable parameters

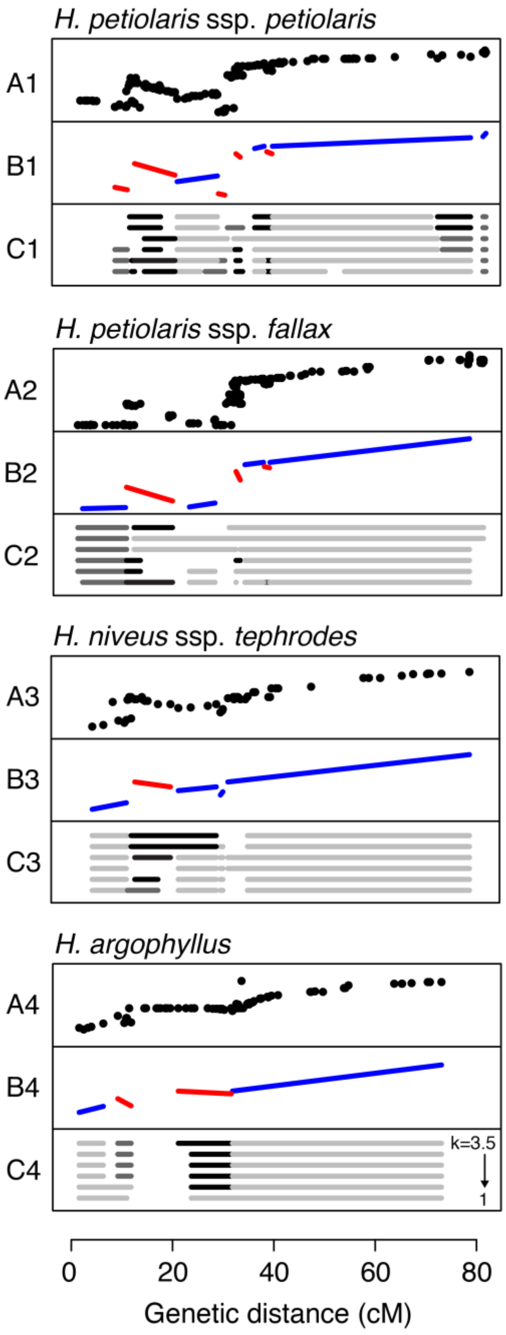

Figure S7 – Example of synteny block assignments from multiple runs of syntR with different $CR_{max}$ parameters and the final curated synteny blocks. A) Marker positions in each individual genetic map relative to the *H. annuus* map on the x-axis. Y-axis is genetic map distance (cM) in each other *Helianthus* taxon. B) The extent of the final curated synteny blocks in both maps and their orientations (blue = positive and red = negative). C) The extent synteny blocks in *H. annuus* found with different max_k_dist parameters and their orientations relative to the other map (starting from the top $CR_{max}$ is 3.5, 3, 2.5, 2, and 1.5, light grey = positive, grey = unclassified, black = negative).

#### Figure S8 - Genetic maps with marker density

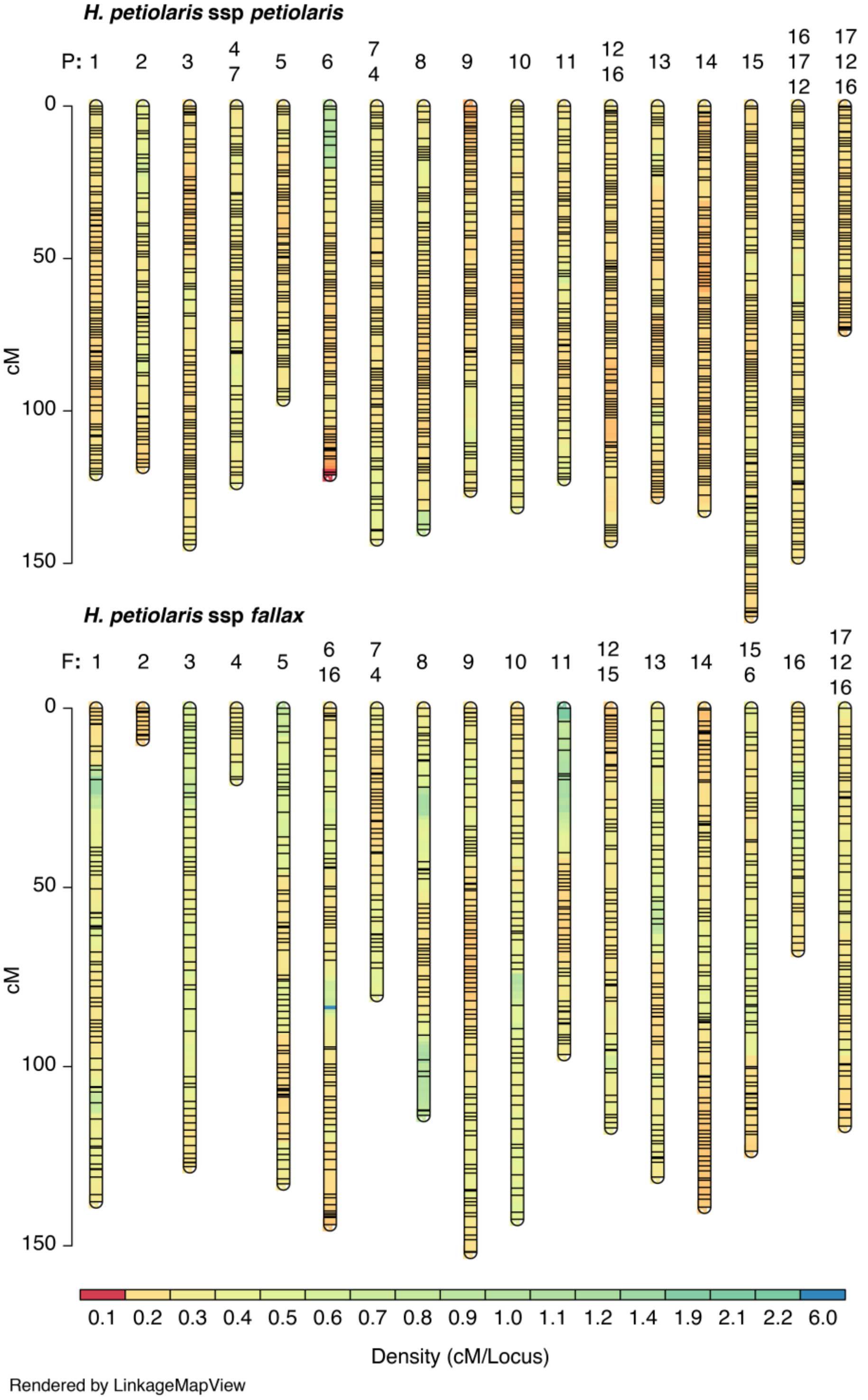

Figure S8 – Marker density of *H. petiolaris* genetic maps. This figure was made with LinkageMapView (Ouellette et al. 2017).

#### Figure S9 – Cumulative marker density plots

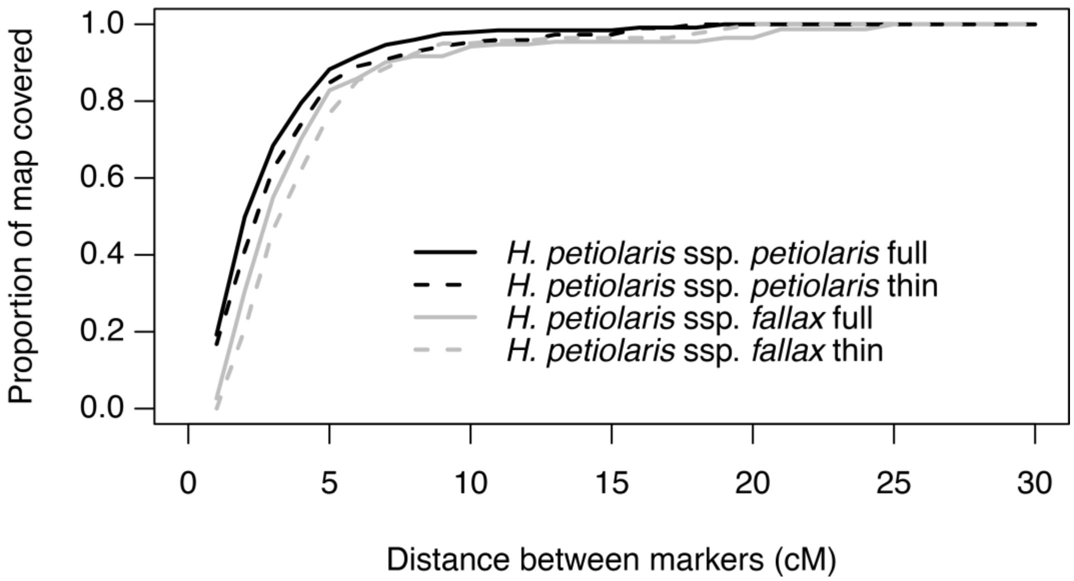

Figure S9 – The extent of *H. petiolaris* genetic maps with a minimum genetic distance between markers.

#### Table S2 – *H. petiolaris* map properties

Table S2 – Properties of new and previously published *H. petiolaris* maps.

| Genetic map | N markers | Length (cM) |
|---|---|---|
| *H. petiolaris* ssp. *petiolaris* - full | 8179 | 1850 |
| *H. petiolaris* spp. *fallax* - full | 13335 | 2178 |
| *H. petiolaris* ssp. *petiolaris* – thinned | 2462 | 1576 |
| *H. petiolaris* spp. *fallax* - thinned | 3368 | 1791 |
| *H. petiolaris* - Burke et al. 2004 | 795 | 1592 |

#### Figure S10 – Full versus thin maps

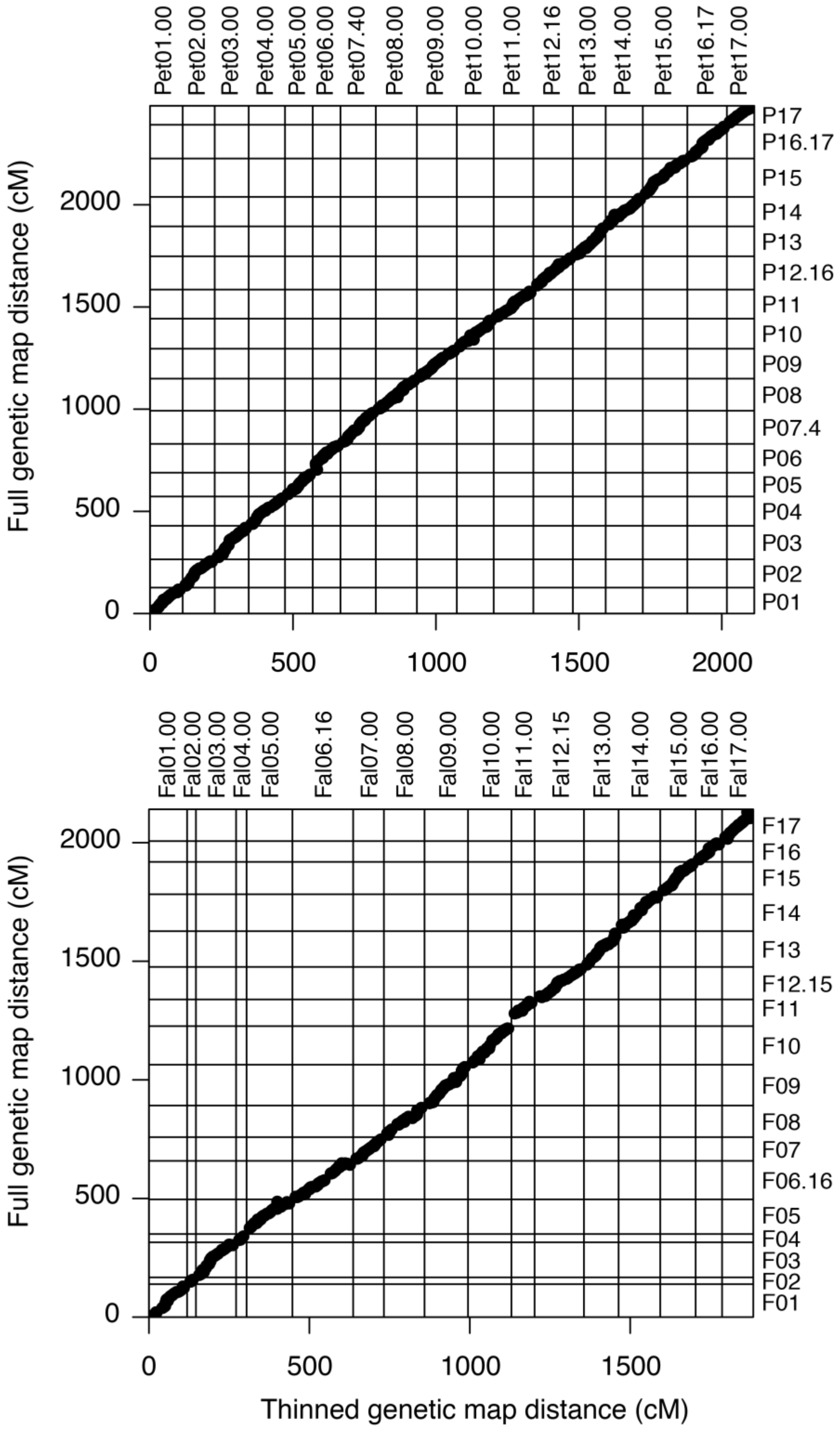

Figure S10 – Comparison of thinned and full *H. petiolaris* genetic maps. Different chromosomes have a 10 cM buffer between them on the x and y axes.

#### Figure S11 – *H. petiolaris* ssp. *fallax* markers

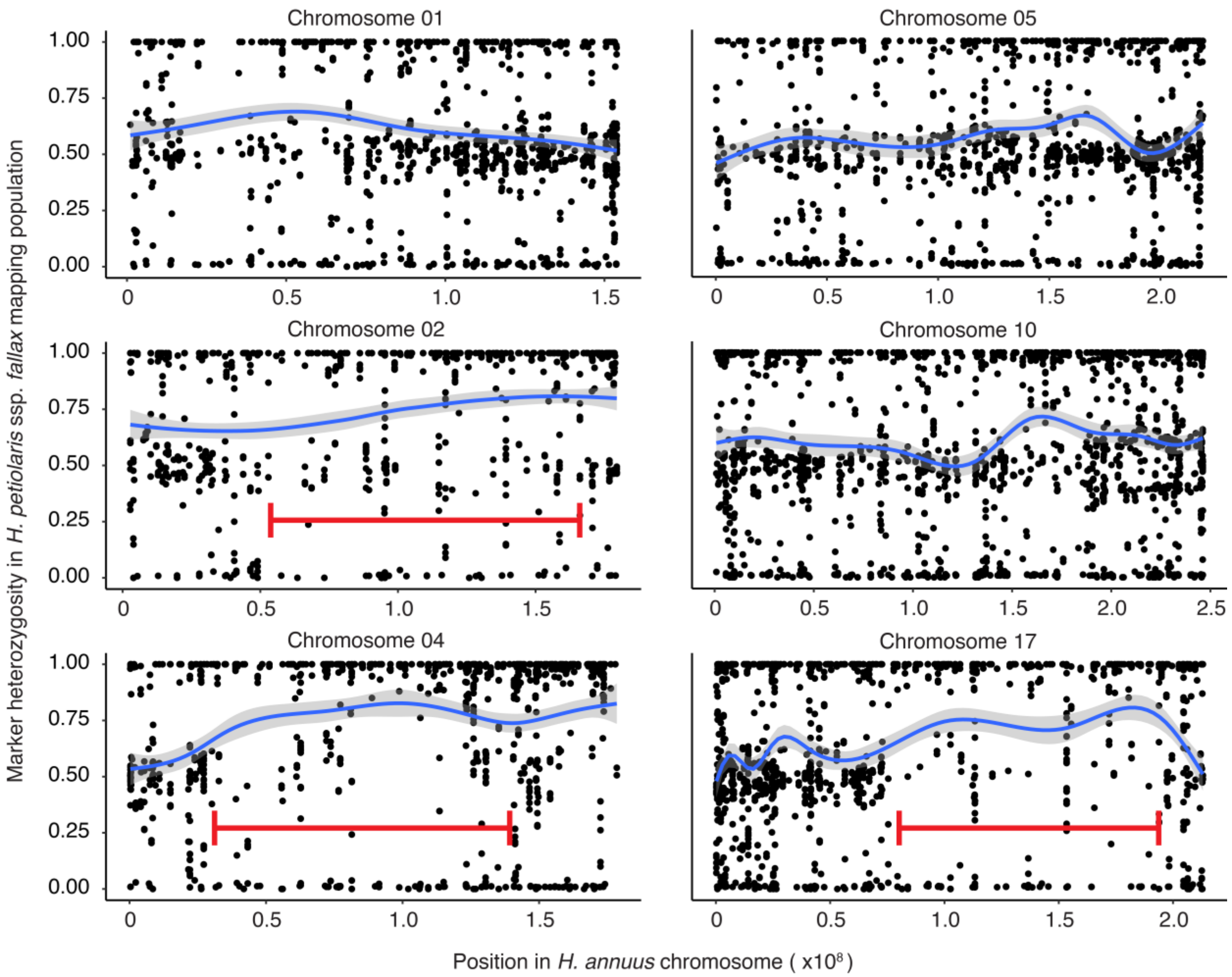

Figure S11 – Heterozygosity of markers in the *H. petiolaris* ssp. *fallax* mapping population relative to position the *H. annuus* reference genome. Blue lines show smoothed average heterozygosity with 95% confidence intervals in grey. Informative markers have heterozygosity values around 0.5. Large sections of *H. annuus* chromosomes 2, 4 and 17 have few markers that are informative (highlighted with red brackets).

#### Figure S12 – *H. petiolaris* maps with synteny blocks

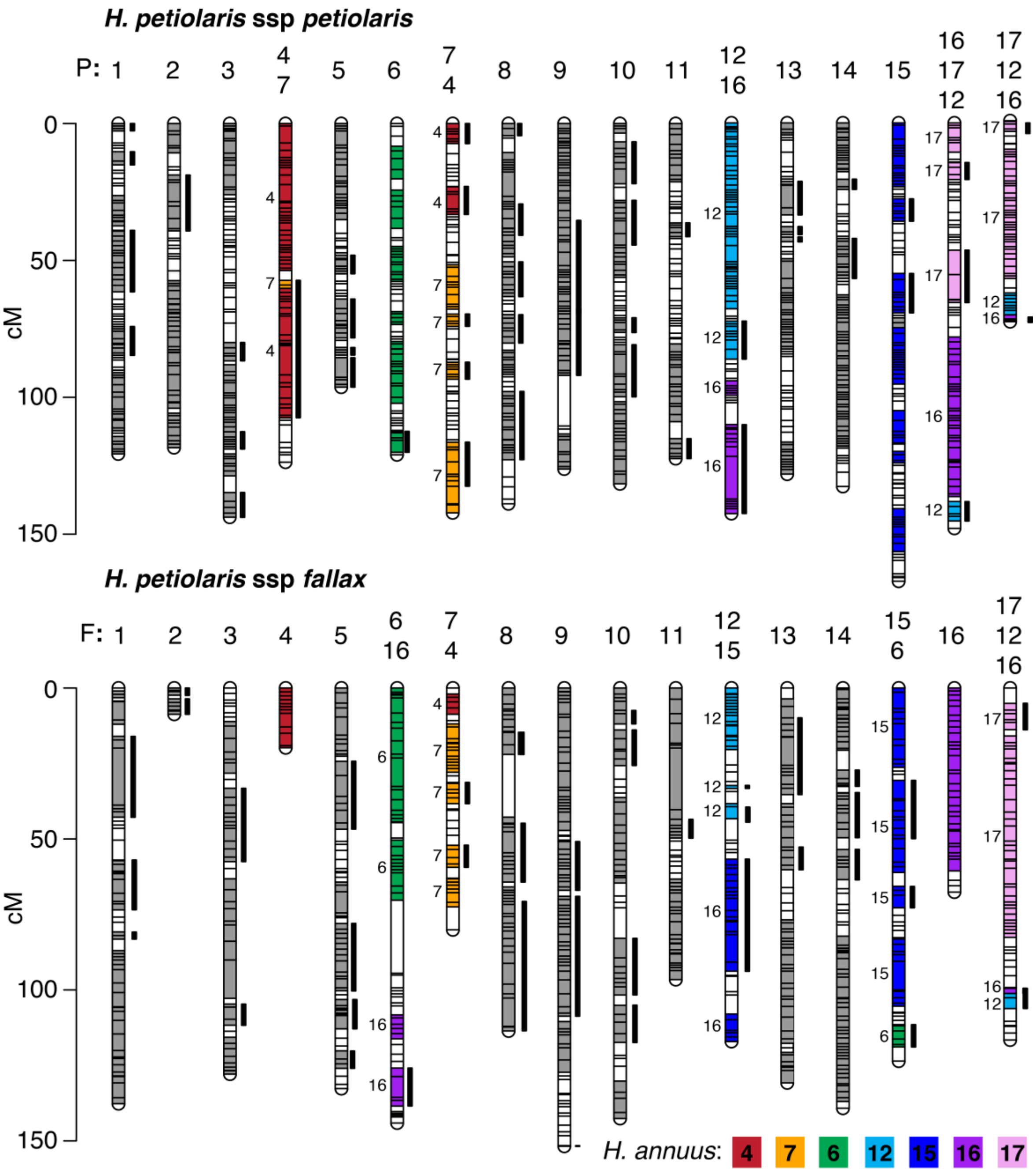

Figure S12 – *Helianthus petiolaris* genetic maps showing blocks of synteny with *H. annuus* found using syntR. Each horizontal bar represents a genetic marker and each thick vertical bar represents a synteny blocks that are inverted relative to the *H. annuus* genetic map. Where there are no translocations between *H. petiolaris* and *H. annuus* chromosomes (e.g. all synteny blocks in P1 and F1 are syntenic with A1), the synteny blocks are shown in grey. Synteny blocks on chromosomes with translocations are color-coded and labeled on the left based on their synteny with *H. annuus* chromosomes. This figure was made with LinkageMapView (Ouellette et al. 2017).

#### Figure S13 – *H. niveus* and *H. argophyllus* maps with synteny blocks

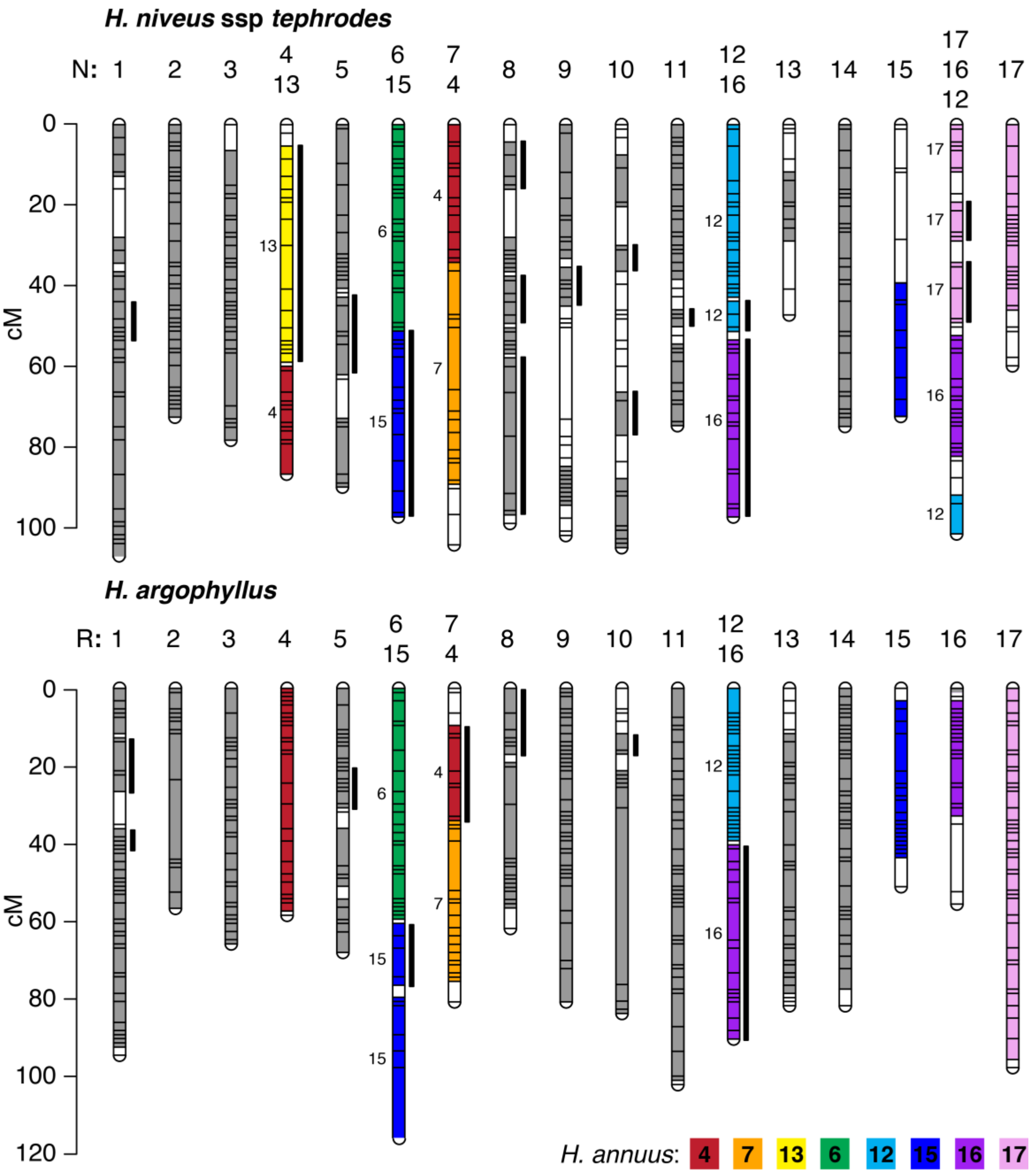

Figure S13 – *Helianthus niveus* ssp. *tephrodes* and *H. argophyllus* genetic maps showing blocks of synteny with *H. annuus* found using syntR. Each horizontal bar represents a genetic marker and each thick vertical bar represents a synteny blocks that are inverted relative to the *H. annuus* genetic map. Where there are no translocations (e.g. all synteny blocks in N1 and R1 are syntenic with A1), the synteny blocks are shown in grey. Synteny blocks on chromosomes with translocations are color-coded and labeled on the left based on their synteny with *H. annuus* chromosomes. The genetic maps are from Barb et al. 2014. This figure was made with LinkageMapView (Ouellette et al. 2017).

#### Figure S14 – Reconstruction of ancestral *Helianthus* karyotypes

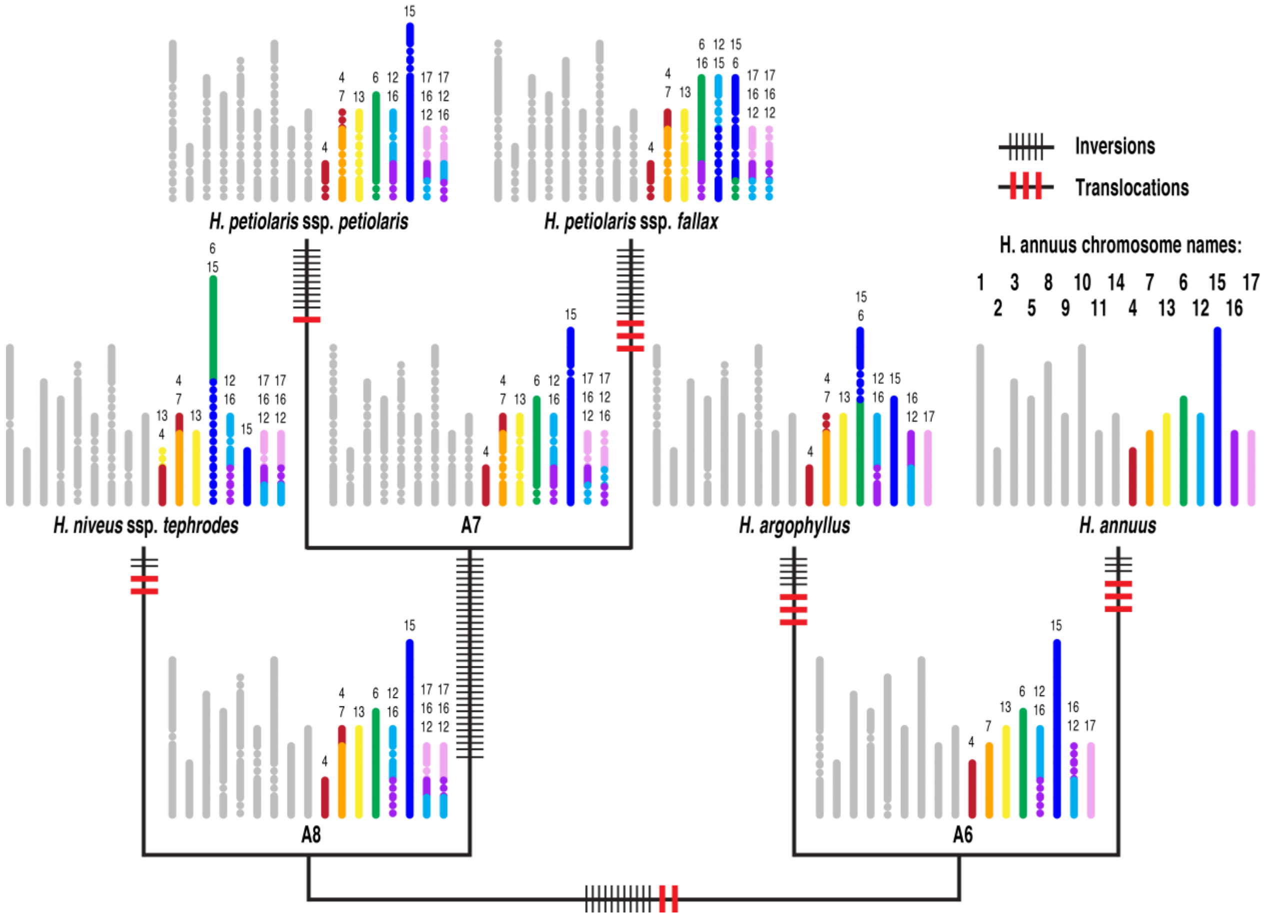

Figure S14 - Diagram of extant and ancestral karyotypes for 5 *Helianthus* taxa. Each synteny block is a fixed size and color-coded based on its position in the *H. annuus* genome (see Fig S21 for a labeled version). Chromosomes without translocations in any many are shown in grey. Inversions are shown using dotted lines. Some synteny blocks were inferred in the additional maps (e.g. part of A17 on F16) when there were some markers supporting it. However, other synteny blocks were dropped (e.g. small region of A7 on P4) because there were no markers to place the block in other maps. Also note that along some branches the same pair of chromosomes is involved in multiple translocations.

#### Table S3 and Table S4 – Divergence time estimates

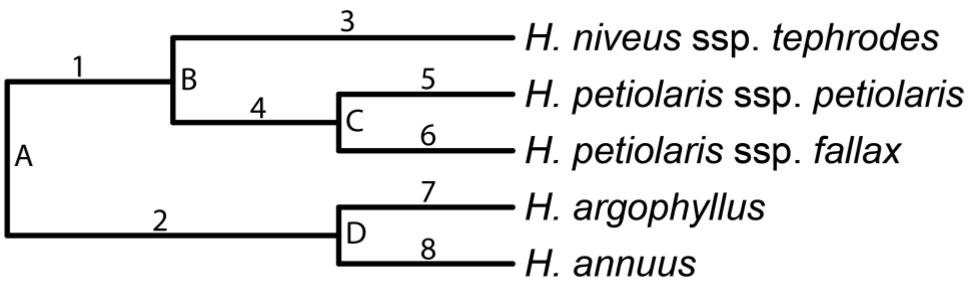

Table S3 – Various divergence time estimates for nodes labeled with letters in the above phylogeny.

| Node | Taxon | Mean | Min | Max | Reference |
|---|---|---|---|---|---|
| A | *H. annuus - H. petiolaris* | 1.8 | | | Sambatti et al. 2012 |
| A | *H. annuus - H. niveus* ssp. *tephrodes* | 1.89 | 0 | 5.25 | Mason 2018 |
| A | *H. annuus - H. petiolaris* | 2.43 | 2.41 | 2.47 | Todesco et al. 2019 |
| C | *H. petiolaris* ssp. *petiolaris - H. petiolaris* ssp. *fallax* | 2.00 | 1.97 | 2.03 | Todesco et al. 2019 |
| D | *H. annuus - H. argophyllus* | 1.54 | 0 | 4.96 | Mason 2018 |
| D | *H. annuus - H. argophyllus* | 1.87 | 1.84 | 1.90 | Todesco et al. 2019 |

Table S4 – Cumulative divergence time estimates based on different various divergence time estimates (Table S3). See phylogeny above to identify the specific numbered branches.

| Branch | Todesco et al. lengths | Todesco et al. min lengths | Todesco et al. max lengths | Max lengths |
|---|---|---|---|---|
| 1 | 0 | 0 | 0 | 0 |
| 2 | 0.56 | 0.57 | 0.57 | 0.29 |
| 3 | 2.43 | 2.41 | 2.47 | 5.25 |
| 4 | 0.43 | 0.44 | 0.44 | 3.22 |
| 5 | 2.00 | 1.97 | 2.03 | 2.03 |
| 6 | 2.00 | 1.97 | 2.03 | 2.03 |
| 7 | 1.87 | 1.84 | 1.90 | 4.96 |
| 8 | 1.87 | 1.84 | 1.90 | 4.96 |
| Total: | 11.16 | 11.04 | 11.34 | 22.74 |

#### Table S5 – Patterns of chromosomal rearrangement

Table S5 – Overall patterns of chromosomal rearrangement inferred based on different sets of synteny blocks. Set 1 synteny blocks are curated based on multiple syntR runs and inferred when missing. Set 2 synteny blocks are curated but present in all five maps. Set 3 synteny blocks are the output from individually optimized syntR runs that are present in all five maps. Rate 1 is the number of rearrangements per million years based on a cumulative divergence time of 11.16 million years (Table S4), while rate 2 is the based on 22.74 million years (Table S4). P-value 1 is the probability of seeing the observed number of inversions and translocations if the rate of inversion was equal to the rate of translocation. P-value 2 is the probability of seeing the observed number of chromosomes involved in translocation if all chromosomes were equally likely to be involved in a reciprocal translocation.

| Synteny blocks | N blocks | N inversions | N trans. | Rate 1 | Rate 2 | N translocated chromosomes | P-value 1: Inversion rate = trans. rate | P-value 2: Trans. rate = across chromosomes |
|---|---|---|---|---|---|---|---|---|
| Set 1 | 97 | 74 | 14 | 7.9 | 3.9 | 8 | $5.1x10^{-11}$ | $8.0 x10^{-8}$ |
| Set 2 | 67 | 45 | 15 | 5.4 | 2.6 | 7 | $1.3 x10^{-4}$ | $5.3 x10^{-11}$ |
| Set 3 | 76 | 50 | 10 | 5.4 | 2.6 | 7 | $1.6x10^{-7}$ | $3.0 x10^{-6}$ |

#### Figure S15 – Non-random translocation probability distributions

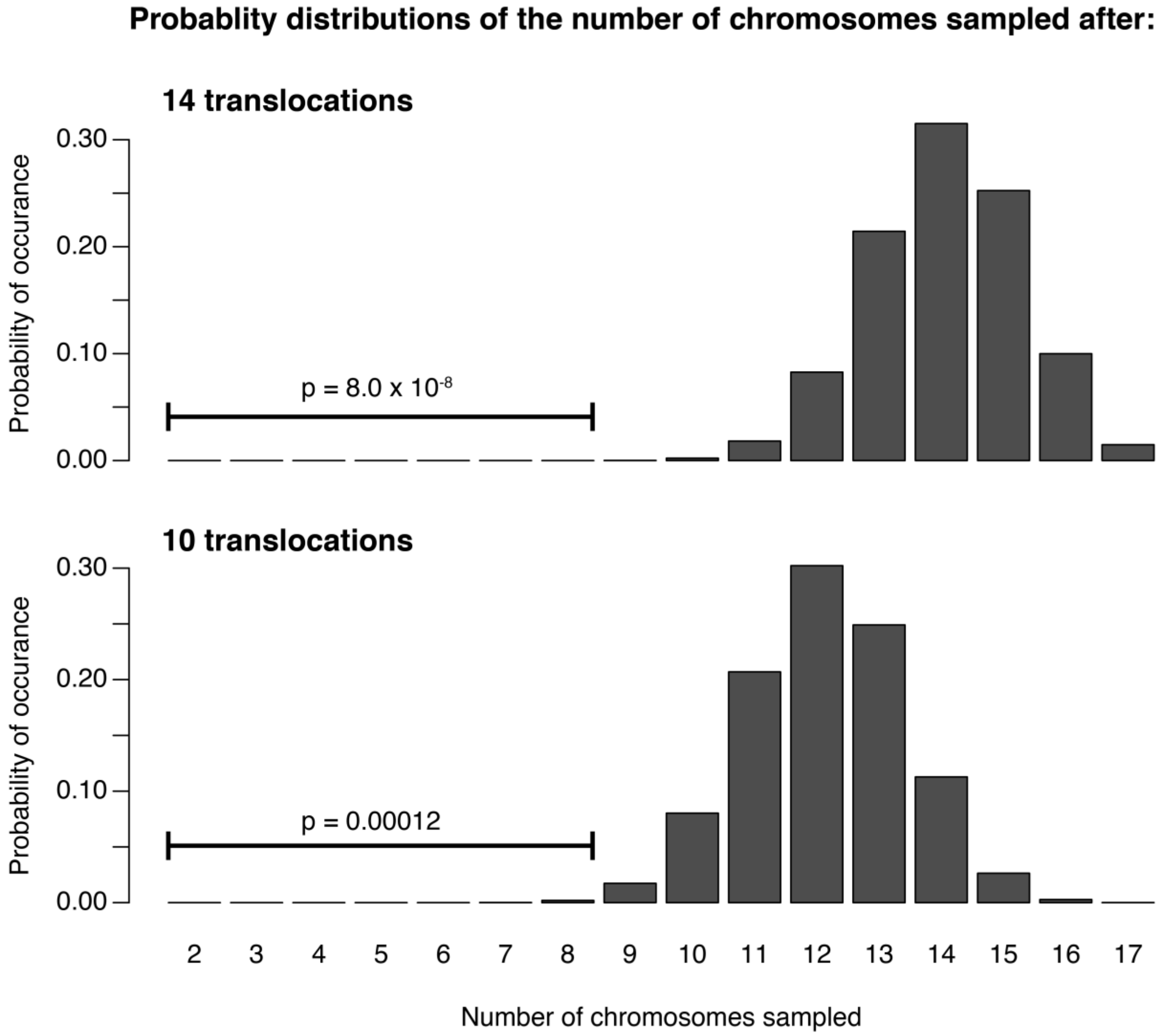

Figure S15 – The probability distribution of sampling some number of 17 possible chromosomes after 14 and 10 translocations. These probability distributions were calculated using random walk with transition probabilities equal to the probability that the number of chromosomes sampled at each translocation remains the same $\frac{\binom{k}{2}}{\binom{17}{2}}$, increases by one $\frac{\binom{17-k}{1}\binom{k}{1}}{\binom{17}{2}}$, or increases by two $\frac{\binom{17-k}{2}}{\binom{17}{2}}$, where k is the number of chromosomes previously sampled. Note that two chromosomes are always sampled during the first translocation.

#### Figure S16 – Chromosome pairing at meiosis

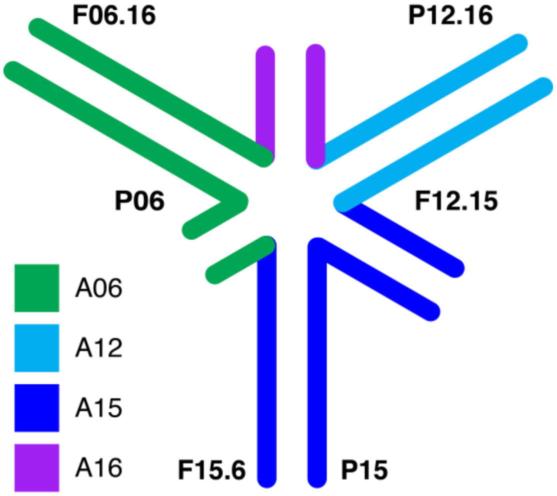

Figure S16 – Hypothetical pairing of chromosomes F06.16, F12.15, F15.16, P12.16, P15, and P06 during meiosis in a hybrid between *H. petiolaris* ssp. *petiolaris* and *H. petiolaris* ssp. *fallax* based on the order and orientation of their synteny blocks. Synteny blocks are color-coded based on their synteny with *H. annuus* chromosomes.

### Figure S17 – *H. petiolaris* ssp. *petiolaris* dot plot with centromere sequence

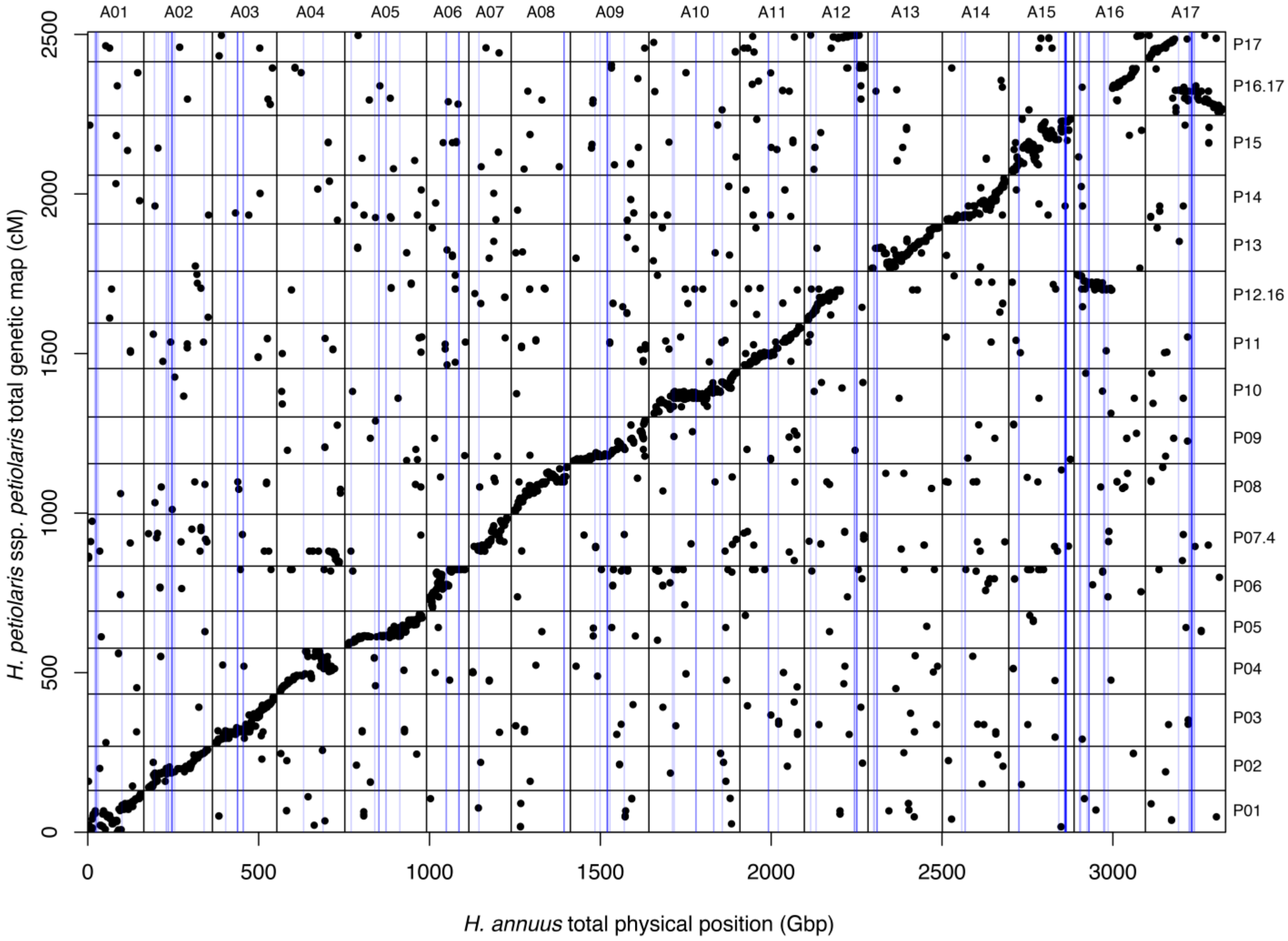

Figure S17 - Dot plot showing the positions of genetic markers the in *H. petiolaris* ssp. *petiolaris* map relative to *H. annuus* physical positions. Chromosomes have a 10 Gbp buffer between them on the x-axis and a 10 cM buffer between them on the y-axis. Blue lines represent the locations in the XRQ sunflower genome with significant similarity (blast hits with evalues < $10^{-5}$) to the centromeric DNA sequence, HaCEN_LINE (Nagaki et al. 2015; Genback accession number: LC075745).

#### Figure S18 – *H. petiolaris* ssp. *fallax* dot plot with centromere sequence

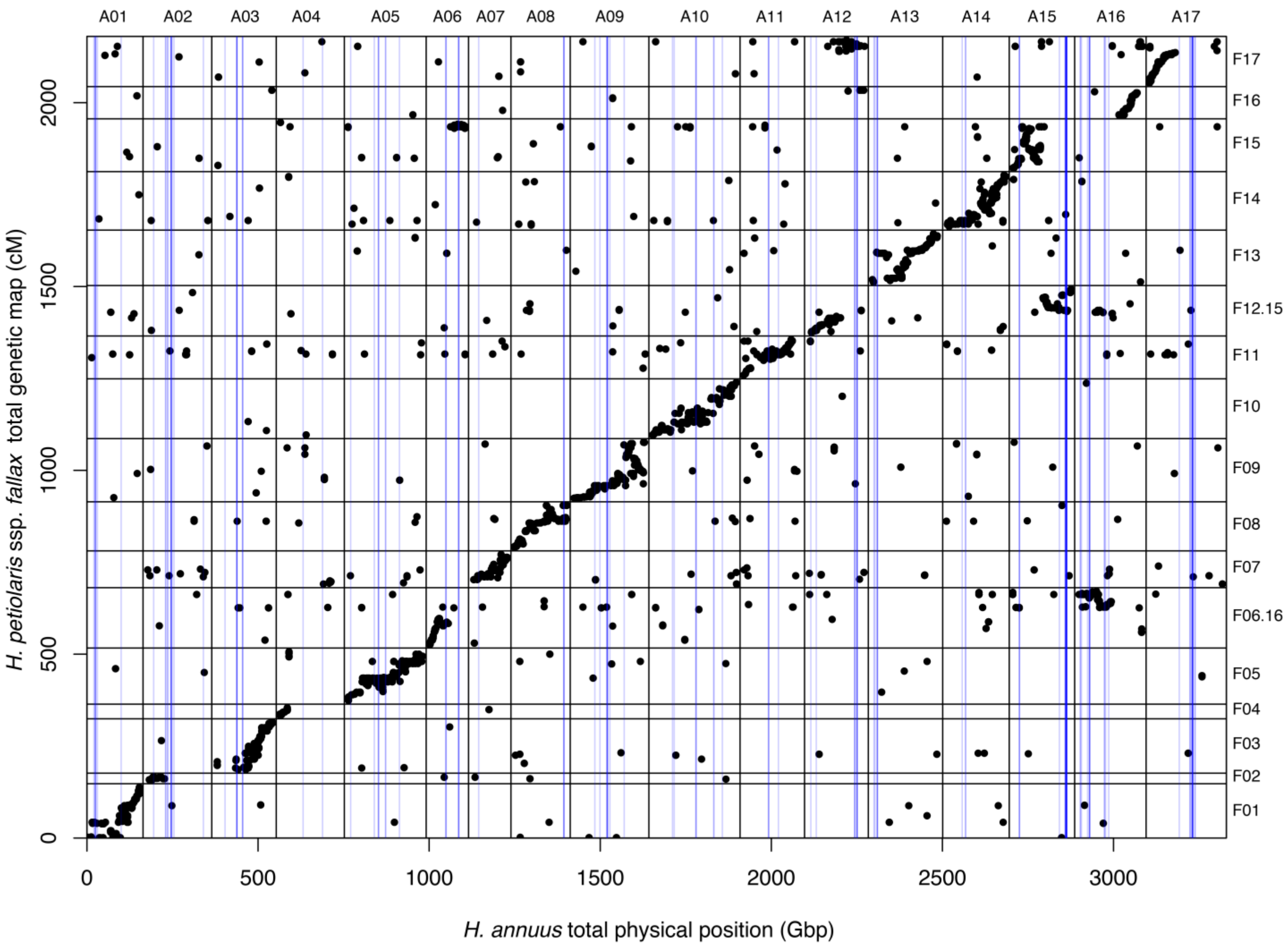

Figure S18 - Dot plot showing the positions of genetic markers the in *H. petiolaris* ssp. *fallax* map relative to *H. annuus* physical positions. Chromosomes have a 10 Gbp buffer between them on the x-axis and a 10 cM buffer between them on the y-axis. Blue lines represent the locations in the XRQ sunflower genome with significant similarity (blast hits with evalues < $10^{-5}$) to the centromeric DNA sequence, HaCEN_LINE (Nagaki et al. 2015; Genback accession number: LC075745).

#### Figure S19 – *H. niveus* ssp. *tephrodes* dot plot with centromere sequence

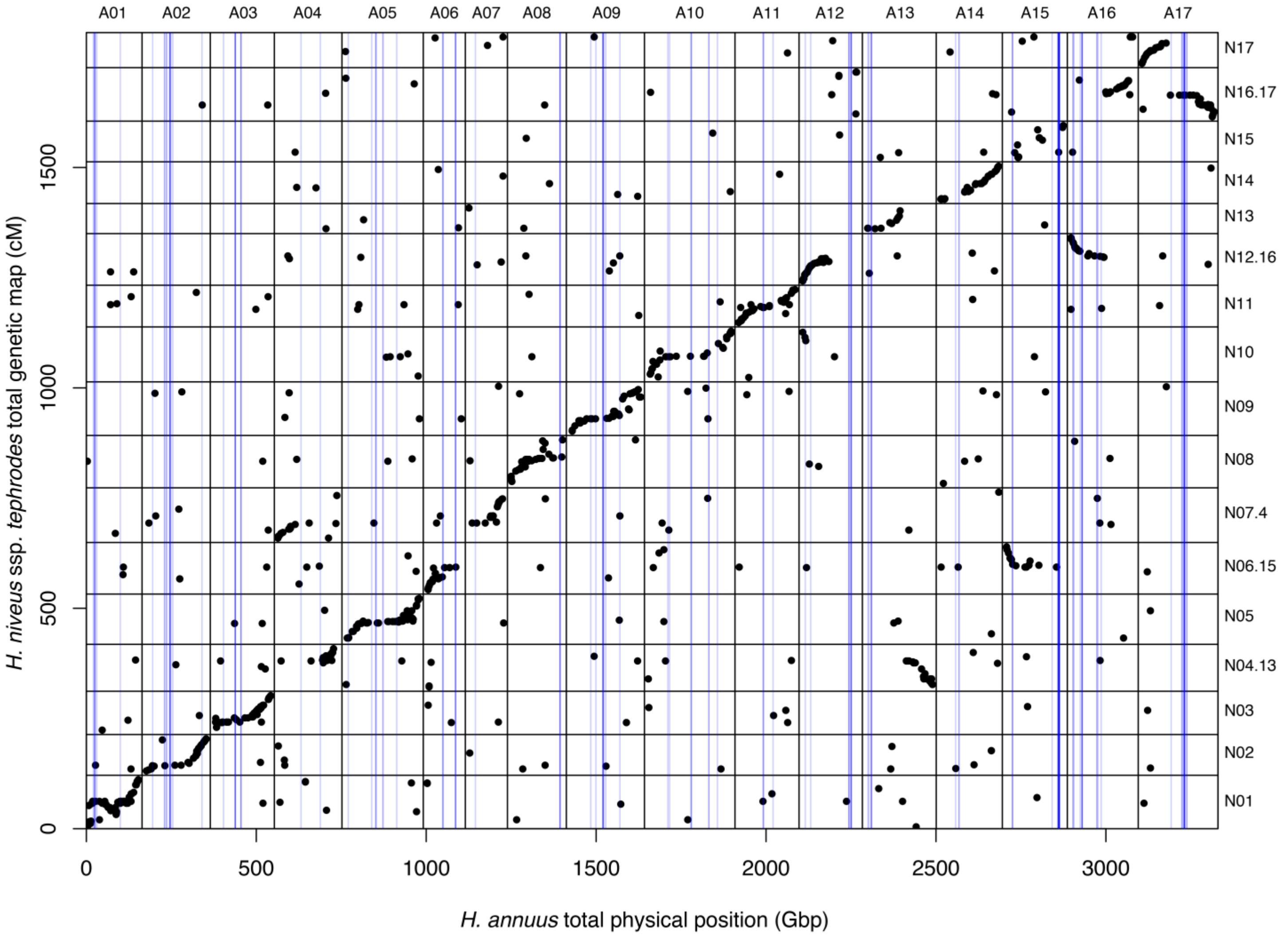

Figure S19 - Dot plot showing the positions of genetic markers the in *H. niveus* ssp. *tephrodes* map relative to *H. annuus* physical positions. Chromosomes have a 10 Gbp buffer between them on the x-axis and a 10 cM buffer between them on the y-axis. Blue lines represent the locations in the XRQ sunflower genome with significant similarity (blast hits with evalues < 10-5) to the centromeric DNA sequence, HaCEN_LINE (Nagaki et al. 2015; Genback accession number: LC075745).

#### Figure S20 – *H. argophyllus* dot plot with centromere sequence

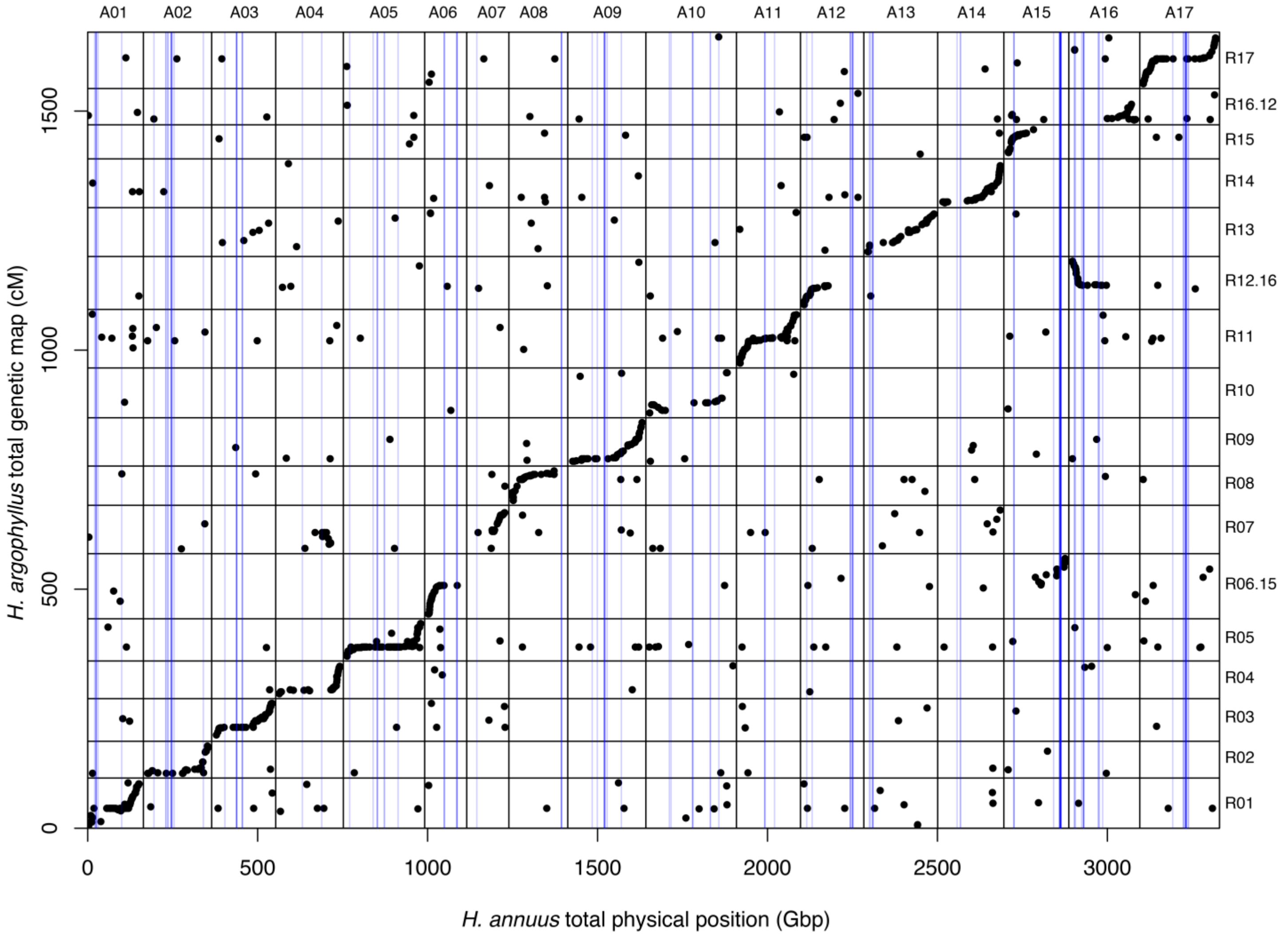

Figure 20 - Dot plot showing the positions of genetic markers the in *H. argophyllus* map relative to *H. annuus* physical positions. Chromosomes have a 10 Gbp buffer between them on the x-axis and a 10 cM buffer between them on the y-axis. Blue lines represent the locations in the XRQ sunflower genome with significant similarity (blast hits with evalues < $10^{-5}$) to the centromeric DNA sequence, HaCEN_LINE (Nagaki et al. 2015; Genback accession number: LC075745).
